# Cell-type-specific architecture of the hypothalamus in a socially plastic vertebrate

**DOI:** 10.64898/2026.06.25.734636

**Authors:** Mélanie Dussenne, Micah Castillo, Preethi H. Gunaratne, Andrew P. Hoadley, Lauren A. Saenz, Beau A. Alward

## Abstract

The hypothalamus orchestrates social behaviors by integrating physiological state with environmental information, but the cellular substrates of this plasticity remain unresolved. We combined single-cell and spatial transcriptomics to generate a cell-type map of the hypothalamus in *Astatotilapia burtoni*, a cichlid fish that forms dynamic social hierarchies. We identified 28 neuronal, glial, neurogenic, and immune cell populations and mapped their organization across hypothalamic nuclei. Social status, sex, and reproductive state engaged coordinated, cell-type-specific transcriptional programs, revealing modular deployment of steroid hormone signaling and plasticity-associated genes. The atlas identified elevated *sst1.1* expression in the hypothalamus of dominant males that we localized to the teleost VMH. CRISPR-Cas9 disruption of *sst1.1* increased body size, suggesting a role for optimal metabolic and energy allocation. These results define a cellular framework for understanding how hypothalamic plasticity enables flexible social behavior.

## Introduction

Social behaviors require cellular systems in the brain that balance stability with flexibility in the face of changing social and physiological conditions. In vertebrates, this balance is coordinated by the hypothalamus, which integrates external cues with endocrine signals to shape context-appropriate behavioral states *(1–4)*. Despite its central role in social and reproductive control, the cellular organization through which the hypothalamus implements behavioral plasticity remains poorly defined.

African cichlids are an experimentally tractable system for addressing this problem. In *Astatotilapia burtoni*, males and females exhibit profound sex differences in reproductive physiology and behavior *(5–8)*. Additionally, males undergo rapid and reversible transitions between dominant (DOM) and subordinate (SUB) social states that differ in aggression, reproductive capacity, and circulating steroid hormone levels (Fig. 1A) *(4, 9)*. These transitions are accompanied by pronounced changes in hypothalamic gene expression and neuroendocrine activity *(10–12)*, allowing cellular analysis of socially induced hypothalamic state.

**Fig. 1.**
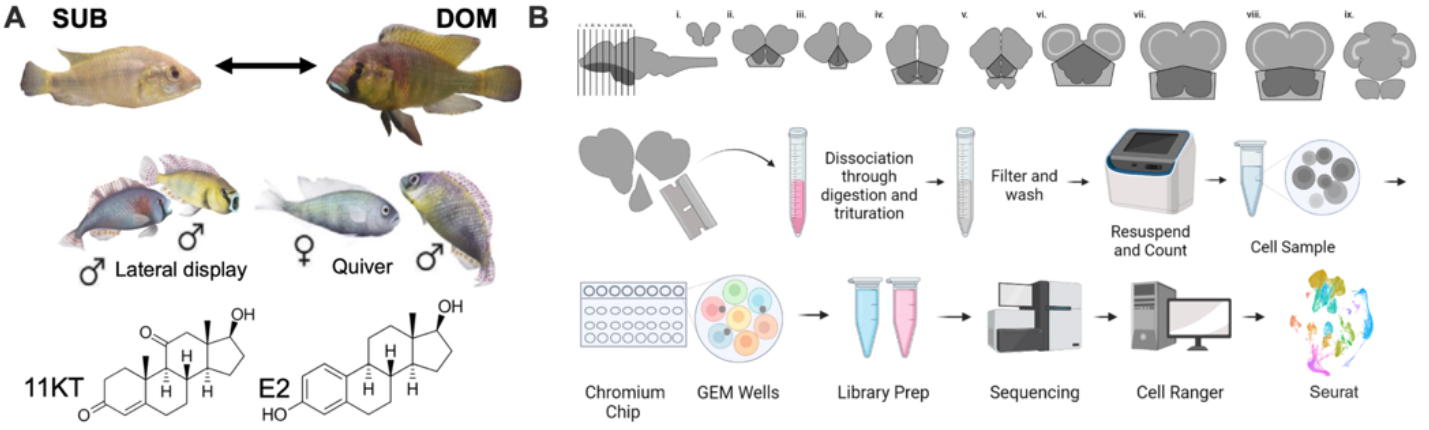
Generating a single-cell atlas of the hypothalamus of a social cichlid. (A) *A. burtoni* males form a social hierarchy with dominant (DOM) and subordinate (SUB) individuals. DOM males have an activated reproductive system and elevated sex steroid hormones 11-ketotestosterone (11KT) and estradiol (E2), and perform territorial displays (lateral display) and court females (quiver). Females exhibit substantial reproductive plasticity: they can be gravid (egg-bearing) or non-gravid. We sample four females and three males spanning reproductive states (gravid, non-gravid, DOM, and SUB) for scRNA-seq. (B) Customized workflow for scRNA-seq in *A. burtoni*. Gray areas denote regions that were sampled.

Recent advances in single-cell and spatial transcriptomic approaches now enable systematic resolution of complex brain regions and their anatomical organization *(13–19)*. Applied to the cichlid hypothalamus, these methods enable the determination of how sex, social status, and reproductive state engage distinct cell populations and molecular programs. Here we combine scRNA-seq and spatial transcriptomics to generate a spatially resolved cell-type atlas. We identify diverse neuronal, glial, and progenitor populations and show that social and physiological variables modulate gene expression in a cell-type–specific manner. Leveraging these findings, we link a somatostatin population in the anterior tuberal nucleus to DOM status. We then show, using CRISPR/Cas9 gene editing, that *sst1.1* inhibits body growth, thereby demonstrating the functional accessibility of our atlas. This *A. burtoni* spatially resolved hypothalamic cell-type atlas provides a cellular blueprint for understanding how social experience shapes hypothalamic function in a socially flexible vertebrate.

## Results

### A molecular atlas of hypothalamic cell types in *A. burtoni*

To define the cellular composition of the *A. burtoni* hypothalamus, we performed scRNA-seq using an optimized dissection and cell isolation workflow to sample the hypothalamus from adult males and females across social and reproductive states (Fig. 1B). Across seven individuals, we obtained high-quality transcriptional profiles from 93,686 cells. Unbiased clustering resolved 28 transcriptionally distinct populations representing neurons, glia, neurogenic progenitors, oligodendrocyte lineage states, and immune cells in males and females (Fig. 2A; 271; Fig. S2). Cell types were annotated using combinations of canonical marker genes and lineage-enriched transcripts, accounting for teleost-specific gene duplications. Several populations were defined by combinatorial expression patterns, reflecting the molecular diversity of hypothalamic cell states in this species.

**Fig. 2.**
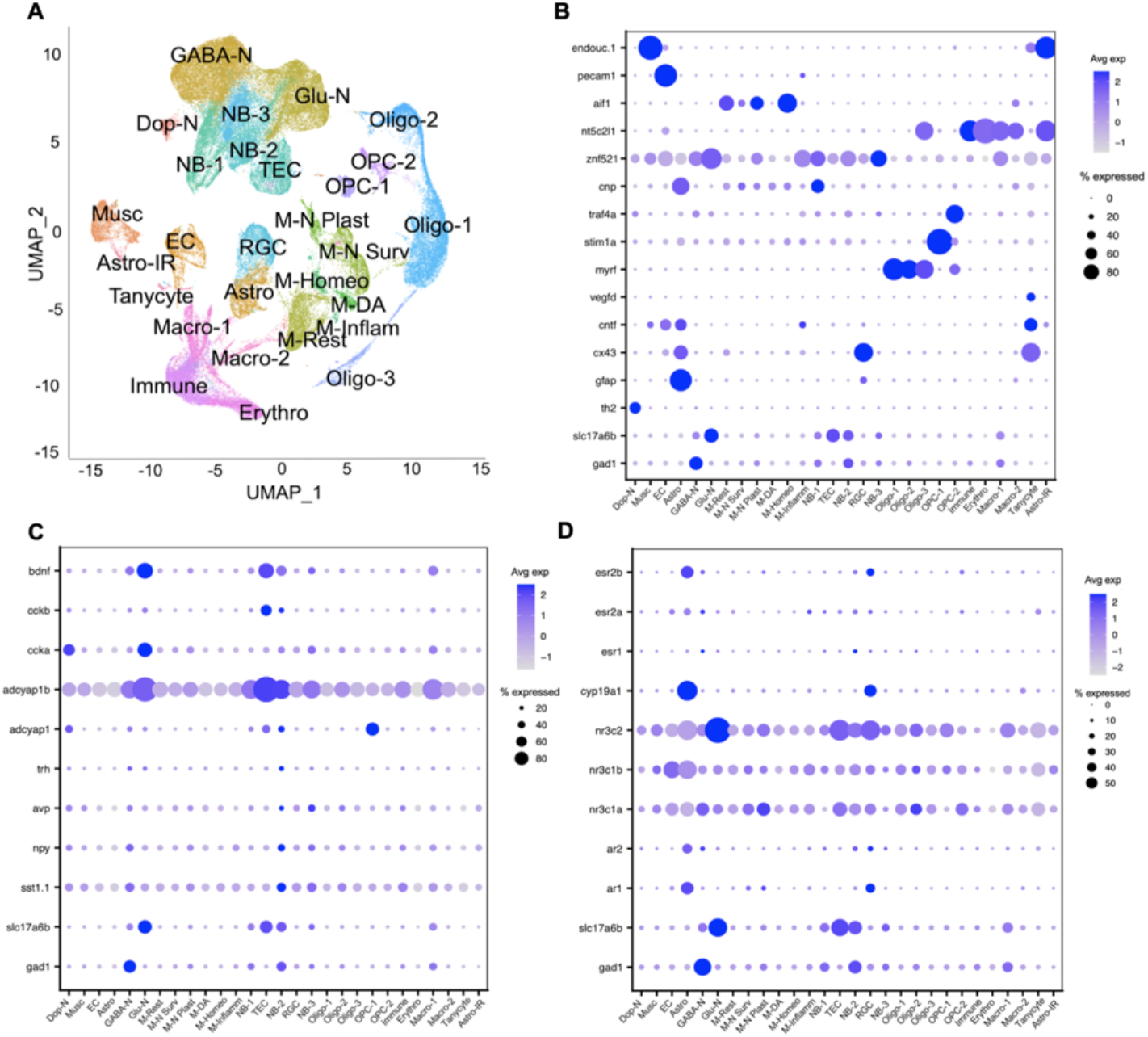
Cell type identification and cell-type specific peptide and steroid gene expression. (A) Uniform manifold approximation and projection (UMAP) showing 28 distinct cell clusters identified using unbiased clustering in Seurat. Cell-type acronyms are overlaid on each distinct cluster. (B-D) Gene expression (y-axis) across cell types (x-axis) is shown as dot plots for (B) marker genes, (C) peptide neuron genes, and (D) steroid signaling genes. N=Neuron; GABA=GABAergic; Glu=glutamatergic; Dop=Dopamine; NB=Neuroblast; TEC=Tanycyte-like ependymal cells; Oligo=Oligodendrocytes; OPC=Oligo precursor cells; Musc=Muscle; EC=Endothelial cells; RGC=Radial glia cells; Astro=Astrocytes; M=microglia; Plast=Plasticity; Surv=Survival; Homeo=Homeostatic; DA=Disease associated; Inflam=inflammatory; Rest=Resting; Macro=Macrophages; Erythro=erythrocytes

We identified glutamatergic, GABAergic, and dopaminergic neurons. Glutamatergic neurons were identified by expression of *slc17a6a* and *slc17a6b*, GABAergic neurons by expression of *gad1*, and a small dopaminergic population by expression of *th2(13)* (Fig. 2B), a teleost-specific tyrosine hydroxylase paralog. A large portion of GABA-N expressed multiple ionotropic glutamate receptor subunit genes, including *grin1a, grin2a*, and *grin2ab* (Fig. S3). Both Glu-N and GABA-N expressed GABA_*B*_ receptor subunit genes *gabbr1a* and *gabbr2*. Thus, hypothalamic GABA-N and Glu-N are sensitive to both inhibitory and excitatory inputs, suggesting integrated excitatory and inhibitory signaling via teleost-specific receptor paralogs.

We identified multiple non-neuronal populations, including tanycytes, radial glial cells (RGC), astrocytes (Astro), oligodendrocytes (Oligo), oligodendrocyte precursor cells (OPC), progenitor and neuroblast cells, and diverse immune cell states. RGC and tanycytes, defined by co-expression of ventricular and glial markers (RGC: e.g., *cx43, ptn*; tanycytes: e.g., *fabp7a, cntf)(20, 21)*, were prominent, consistent with the presence of persistent neurogenic niches in the adult teleost brain. A small subset of tanycytes highly expressed *vegfd*, a marker typically associated with lymphatic cells *(22, 23)*. Canonical neuroblast (NB) markers identified three NB clusters: NB1 and NB2 expressed high levels of *stmn2a(24–26)*, whereas NB3 was defined by *znf521* expression *(27–29)*. NB clusters were further distinguished by *cnp* (NB1) and *rbms3* (NB3) expression. Another cluster was identified as tanycyte-like ependymal cells (TEC) based on expression of genes involved in epithelial tight junctions, secretory barriers, and cell adhesion, including *plekha7, pard6gb, aqp3a.1, slc27a1a*, and *rhbg(30*–*50)*. In mice, hypothalamic TEC can function as neural progenitors and are modulated by afferent neural activity *(51–55)*. Supporting this, TEC highly expressed *gabbr1a, gabbr2, grin1a*, and *grin2ab* (fig. S3), suggesting they are sensitive to neural input that could regulate proliferation.

Immune populations exhibited transcriptional diversity, including macrophages and several microglial states associated with resting-state *(p2ry12)*, homeostasis *(vsir)*, plasticity *(edaradd, ccl20)*, and inflammatory/disease responses *(rps29, nrros, fgl2a, f13a1b, thbs1b)* (fig. S4; fig. S5) *(56–74)*. Together, these data reveal a diverse cellular landscape in the cichlid hypothalamus.

We further analyzed the clusters for peptidergic cell types known to be GABAergic or glutamatergic *(13)*. GABA-N cells highly expressed *npy, sst1.1, adcyap1b*, and *bdnf*, whereas Glut-N cells highly expressed *adcyap1b, ccka*, and *bdnf* (Fig. 2C; Fig. S6A). About 20-50% of Dop-N cells expressed low to moderate levels of *adcyap1b, adcyap1*, and *ccka*. A subset of NB2 cells displayed mature GABA-N characteristics, including expression of *npy, sst1.1, adcyap1b, cckb*, and *bdnf*, suggesting they may represent mature GABA-N retaining neuroblast markers or a proliferative GABA-N subtype. TEC cells expressed the glutamatergic marker *slc17a6b* along with *adcyap1b, cckb*, and *bdnf*, suggesting differentiation toward glutamatergic identities while retaining TEC markers. These patterns may reflect specialized neurogenic states, potentially related to the presence of teleost-specific paralogs of hypothalamic genes such as *adcyap1b* and *cckb*. Thus, although hypothalamic peptidergic GABA-N and Glut-N resemble those in mice *(13), A. burtoni* may possess unique neural and neurogenic cell states that may contribute to its robust adult social and reproductive plasticity.

Recent mouse scRNA-seq studies have shown that steroid signaling genes are expressed in GABAergic and glutamatergic neurons in the hypothalamus *(13)*. We mapped the expression of glucocorticoid receptor genes *(nr3c1a, nr3c1b)*, the mineralocorticoid receptor gene *nr3c2*, and sex steroid signaling genes *ar1, ar2, pgr, esr1, esr2a, esr2b*, and *cyp19a1*, which encodes brain-specific aromatase in teleosts *(5)*, across cell types. *Nr3c1a, nr3c1b*, and *nr3c2* were broadly expressed across hypothalamic cell types, with nearly 100% of Glu-N expressing very high levels of *nr3c2* (Fig. 2D; Fig. S6B). In contrast, sex steroid signaling genes showed greater specialization: Astro, specific OPC and NB populations, and distinct MG states exhibited unique *ar* and *esr* expression patterns. Astro and RGC showed similarly high expression of sex steroid signaling genes, including *ar1, ar2*, and *esr2b*, with distinctly high enrichment for *cyp19a1*. These findings suggest that steroid modulation of hypothalamic function is partitioned among discrete cellular modules in *A. burtoni* and may differ substantially from that in mice.

### Sex-, status-, and reproductive-state–dependent modulation of gene expression

*A. burtoni* exhibit robust sex-, status-, and reproductive-state-dependent differences in social behavior, neuroplasticity, and physiology *(4, 6, 8)*. We discovered pronounced sex differences in cell-type-specific gene expression (Fig. S7, S8). For example, in GABAergic neurons alone, more than 600 genes were differentially expressed between males and females. Female-biased genes included axonal growth inhibitor genes such as *rtn4rl2a* and *rtn4rl2b*. In contrast, males showed elevated expression of genes associated with neurogenesis and metabolic regulation, suggesting sex differences in the transcriptional programs that underlie structural plasticity. Cell-type-specific gene ontology (GO) analysis revealed significant biological process terms across three cell types (Fig. S9). Oligo2 were enriched for cell and synaptic signaling terms, and NB3 were enriched for morphogenesis genes. Microglia involved in neuroplasticity were enriched for trans-synaptic signaling and amide biosynthetic processes, consistent with microglial roles in shaping sex differences in the brain via amidergic mechanisms (e.g., anandamide) *(75)*.

Social status similarly shaped transcriptional programs. DOM males showed increased expression of neurogenic regulators *(emx3, nrxn3a)* and neuropeptides *(avp, sst1.1, npy)* across neurogenic and neuronal populations (Fig. 3A; Fig. S10). SUB males upregulated genes associated with synaptic and circuit formation and stabilization, including *cbln2b* and *robo2*. GO analyses identified enrichment for other biological and molecular processes that differentiate DOM and SUB (Fig. S11). NB GO enrichment highlighted gene programs related to the development of the central, autonomic, enteric, and peripheral nervous systems, as well as neuropeptide signaling and neuronal differentiation (NB1, NB2). In parallel, strong enrichment for neuron projection morphogenesis, axogenesis, dendrite development, and cell–cell adhesion is consistent with circuit remodeling and membrane specialization. At the same time, recurrent epithelial and endocrine differentiation terms suggest persistent engagement of neuroendocrine lineage programs across NB populations. Tanycyte GO terms were enriched for lipid biosynthetic processes and prostaglandin synthesis. Both are thought to be involved in shaping neuroendocrine cellular function, including the plasticity of GnRH-1 neurons, which was recently shown to be controlled by glial prostaglandins in mice *(76)*. These differences suggest distinct neuroplasticity-associated cellular states between DOM and SUB males.

**Fig. 3.**
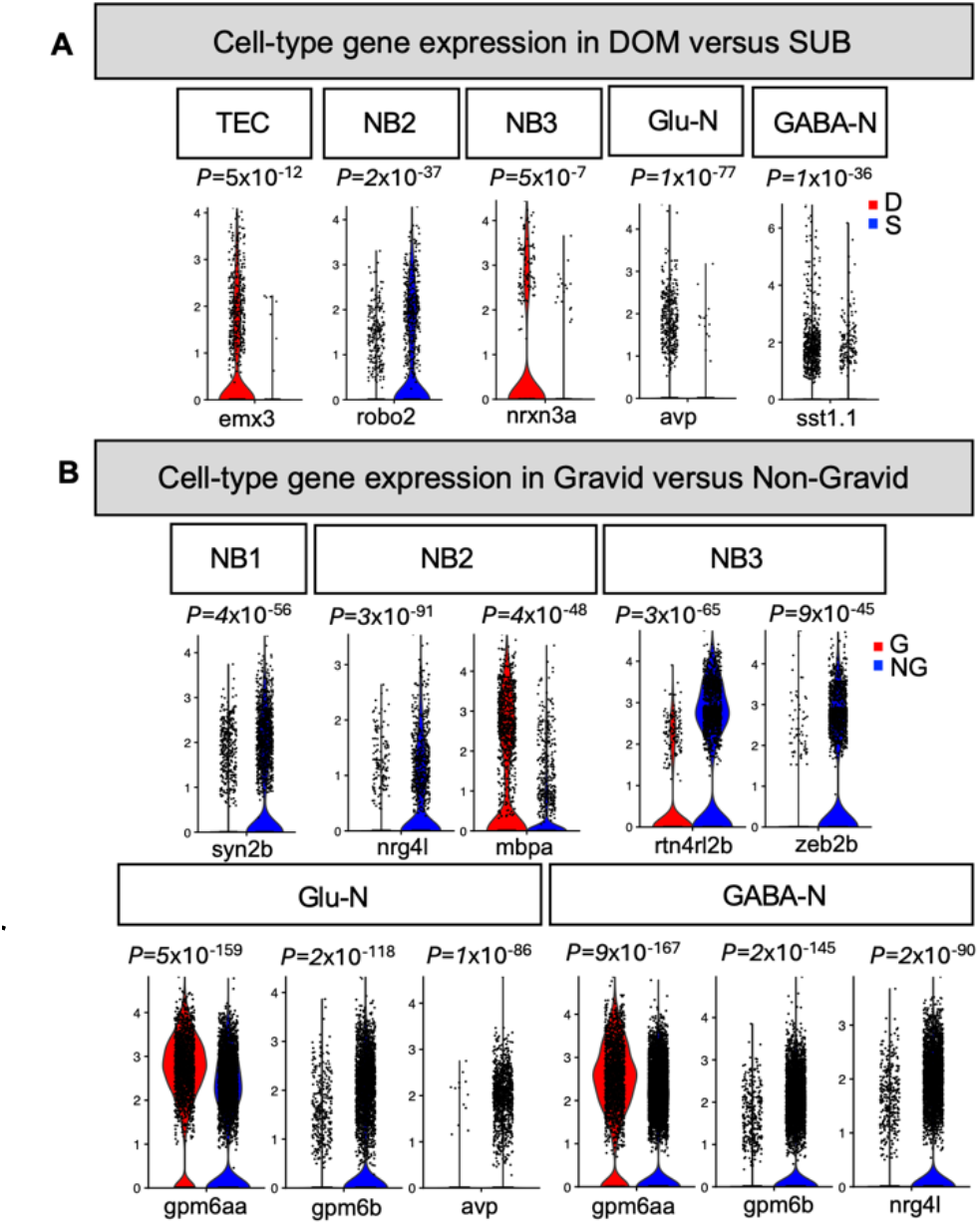
Plasticity and peptide gene expression differ based on cell type, status, and reproductive state. Violin plots show gene expression in specific cell types, comparing (A) DOM versus SUB and (B) Gravid versus Non-Gravid. (A-B) The y-axis shows the expression level. Each data point represents a cell. Adjusted p-values are shown directly above each violin plot. The gene for which expression is shown in each violin plot is on the x-axis.

Female reproductive state also modulated gene expression. Gene programs across virtually all cell types exhibited extensive differences between gravid and non-gravid females, suggesting substantial cellular and circuit remodeling underlying changes in reproductive physiology (Fig. 3B; Fig. S12). Non-gravid females exhibited higher expression of numerous genes involved in neural circuit refinement and neurogenesis, including *syn2b, nrg4l, rtn4rl2b*, and *zeb2b. Mbpa* expression was higher in gravid NB2 cells, indicating gravid females may possess a subset of NB2 cells that were beginning to differentiate to oligodendrocytes. Two *gpm6* genes—*gpm6aa* and *gpm6b—*were higher and lower in Glu-N and GABA-N in gravid females compared to non-gravid females, respectively. *Gpm6a* was shown in mammalian models to enhance neuronal migration, differentiation, and synaptogenesis *(77, 78)*. Recent work suggests that *gpm6b* is required for gliogenesis and neurogenesis *(79)*, while another study suggests it is necessary for the homeostatic regulation of cell differentiation *(80)*. *Nrg4l* was expressed at higher levels in non-gravid GABA-N, mirroring the pattern observed for NB cells. Brain-derived *Nrg4* is not known to play a role in adult brain plasticity (instead, neuregulin-4 released from adipose tissue affects metabolic functions). On the contrary, *Nrg4* is thought to play a pivotal role in shaping brain circuitry during early development, suggesting a potential role for *nrg4l* in adult hypothalamic plasticity in female *A. burtoni*. Immune, neurogenic, and glial cells showed enrichment for GO terms related to cellular adaptation, synaptic signaling, behavior, and structural remodeling (Fig. S13). Altogether, these findings demonstrate substantial cell-type-specific differences in gene programs in *A. burtoni* as a function of sex, status, and reproductive state, and provide a framework for future functional studies of adult brain plasticity.

### Spatial resolution of genetically defined hypothalamic cell populations

We used spatial transcriptomics (Xenium, 10x Genomics) to map scRNA-seq-defined cell populations onto the adult hypothalamus (Fig. 4). A custom 50-gene panel included 35 scRNA-seq-derived gene markers, nine steroid-related (ST) genes, and six neuropeptide (PEP) genes implicated in socio-sexual behavior, including *avt, gnrh1*, and *sst1.1(81–84)*. We also included *npy*, since its expression varies with sex and reproductive state and is co-expressed with *sst1.1* across vertebrates *(85–88)*, as well as the activity-dependent gene *egr1* (Table S1). 35 marker genes were used for clustering and spatial resolution, whereas the remaining 15 genes were projected onto defined cell populations to resolve their spatial organization.

**Fig. 4.**
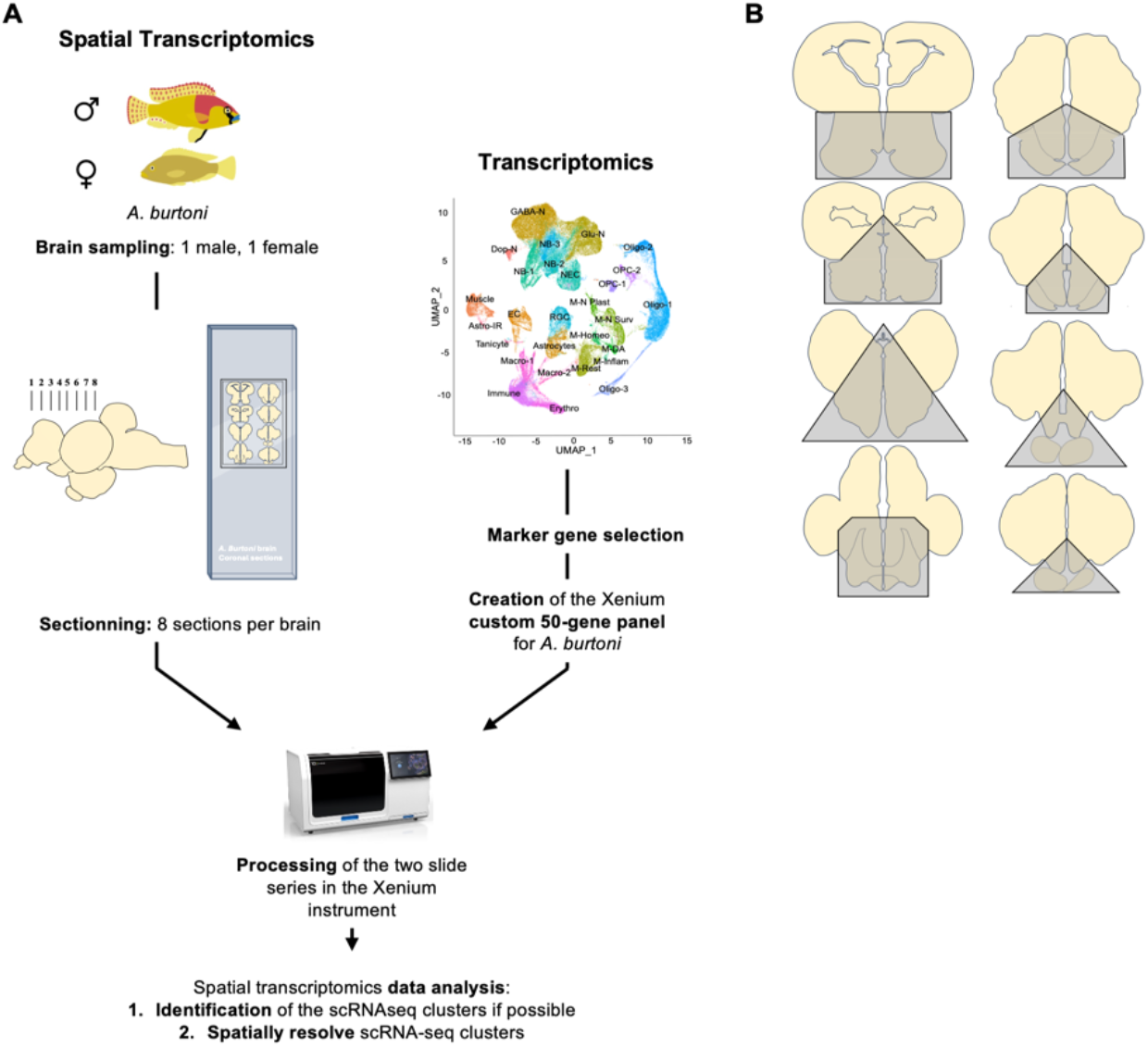
Spatial resolution of hypothalamic cell types using spatial transcriptomics. (A) Representation of the Xenium workflow. One male and one female brain were sampled for spatial transcriptomics analysis. We created a custom Xenium gene panel based on scRNA-seq data. This gene panel was used to process brain slides on the Xenium instrument to spatially resolve transcriptomic cell clusters. (B) Grey areas represent hypothalamic regions selected from brain sections processed for spatial transcriptomics and correspond to the regions sampled for the scRNA-seq analyses. Only these regions were further analyzed.

We profiled one ascending male and one pre-gravid female across 16 hypothalamic sections processed simultaneously by the Xenium instrument (Fig. 4A). After restricting analyses to scRNA-seq-matched regions (Fig. 4B), ~200,000 hypothalamic cells remained for analysis. Unsupervised clustering recovered 27 male and 26 female clusters with highly similar transcriptional and spatial organization (Fig. S14).

Most major scRNA-seq-defined cell classes were spatially resolved. However, dopamine neurons, neuroblast subtypes, macrophages, and some microglial subclasses were not recovered, likely due to reduced gene coverage and tissue sampling (Fig. 5). Glutamatergic neurons occupied discrete nuclei, including the PGc, habenulae, PGl, and TL. In contrast, GABAergic neurons were broadly distributed throughout the hypothalamus *(89)*. Additional populations lacking canonical glutamatergic or GABAergic markers instead expressed glutamate and/or GABA receptor subunits, suggesting anatomically restricted neurotransmitter-responsive states enriched in posterior hypothalamic regions.

**Fig. 5.**
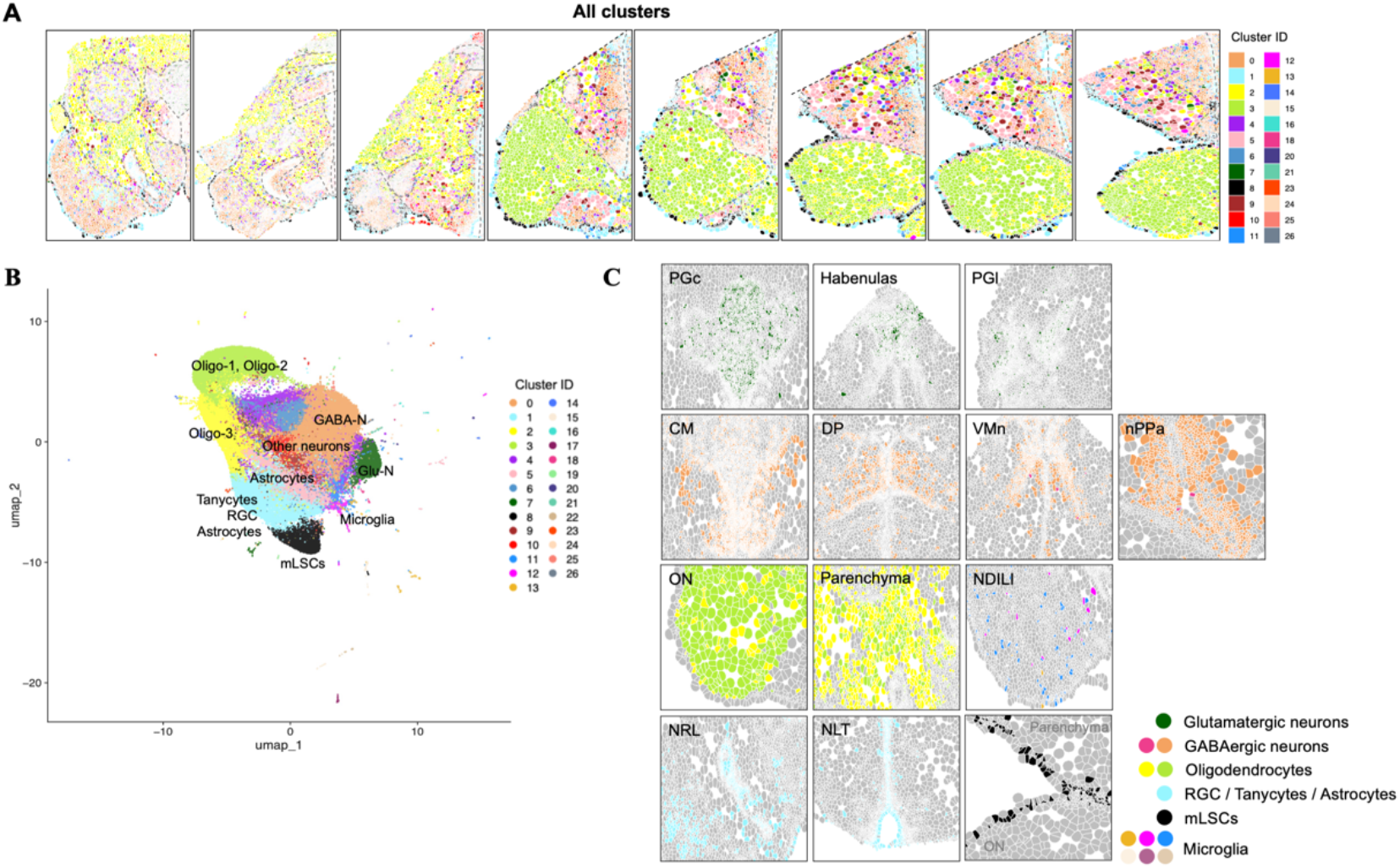
Topography of major hypothalamic cell types identified by spatial transcriptomics. (A) Spatial distribution of all transcriptionally defined cell types across the male *A. burtoni* hypothalamus, from the posterior (left) to the anterior hypothalamus (right). Individual cells are outlined in grey and colored according to their cluster assignment. Dashed lines delineate major hypothalamic nuclei. (B) UMAP represents all profiled hypothalamic cells. (C) Spatial localization of each major cell type. Cells are colored by cluster identity. Abbreviations correspond to the hypothalamic nuclei shown in the panel (Table S2).

Glial populations showed strong spatial segregation. A mixed astrocyte/tanycyte/RGC population lined the third ventricle and pial surface (Fig. 5C). Oligodendrocytes localized mainly outside hypothalamic nuclei and within the optic nerve. In contrast, OPCs were broadly distributed throughout hypothalamic tissue. One large pial population likely represented meningeal lymphatic supporting cells (mLSCs). Microglia were dispersed throughout the hypothalamus, particularly in posterior nuclei.

### Definition of cell types

#### Glial cells

Oligodendrocytes segregated into two major populations distinguished primarily by *mbpa, myrf*, and *pip5kl1* expression (Fig. S15). Oligo-1 cells expressed *mbpa* and *myrf* and were broadly distributed throughout the parenchyma and optic nerve, whereas Oligo-3 cells showed strong *mbpa* expression with little or no *myrf* or *pip5kl1*. Both populations contained subsets expressing *cyp19a1b* and *nr3c2*, indicating the capacity for estradiol synthesis and corticosteroid sensitivity, consistent with observations in humans *(90)* (Fig. S16). OPCs 1 and 2 were identified by *stim1a* or *traf4a* expression. OPC-1 cells were rare and detected only in the male, whereas OPC-2-like populations were transcriptionally heterogeneous and intermixed with neuronal and oligodendrocyte markers (Fig. S17).

A major ventricular and pial cluster expressed *fabp7a, cx43, gfap*, and *rps29*, consistent with radial glia, tanycytes, and astroglia populations (Figs. S18, S19). These cells were present in the NRL, lined the ventricular zones and the outer brain surface, and displayed extensive co-expression of *cyp19a1b* and *nr3c2*, supporting the role of radial glia as major steroid-responsive and estrogen-synthesizing cells in the *A. burtoni* hypothalamus *(91, 92)*. Additional *gfap*+ astroglial populations were distributed throughout the parenchyma, particularly in posterior hypothalamic nuclei, although limited marker resolution prevented complete separation from neurons and oligodendrocytes.

A distinct *vegfd+/cx43+/fabp7a+* population formed a continuous layer surrounding the brain and optic nerve (Fig. 6). Based on their transcriptional profile and localization, these cells likely represent meningeal lymphatic supporting cells (mLSCs) and/or radial astrocytes, recently described in zebrafish (95, 96). The pia mater contains meningeal lymphatic cells, including muLECs (mural lymphatic endothelial cells), the development of which depends on the production of *vegfc* and/or *vegfd* by their primary producers, Radial astrocytes (RAs) *(93–96)* and mLSCs (meningeal lymphatic supporting cells) *(94)*. These RAs were shown to express *gfap* and *cx43*, while both RAs and mLSCs express high levels of *vegfd(93–96)*. Given their location, the cells we observe lining the whole brain are likely mLSCs / RAs. Subsets of these putative mLSCs/RAs expressed *cyp19a1b* and *nr3c2*.

**Fig. 6.**
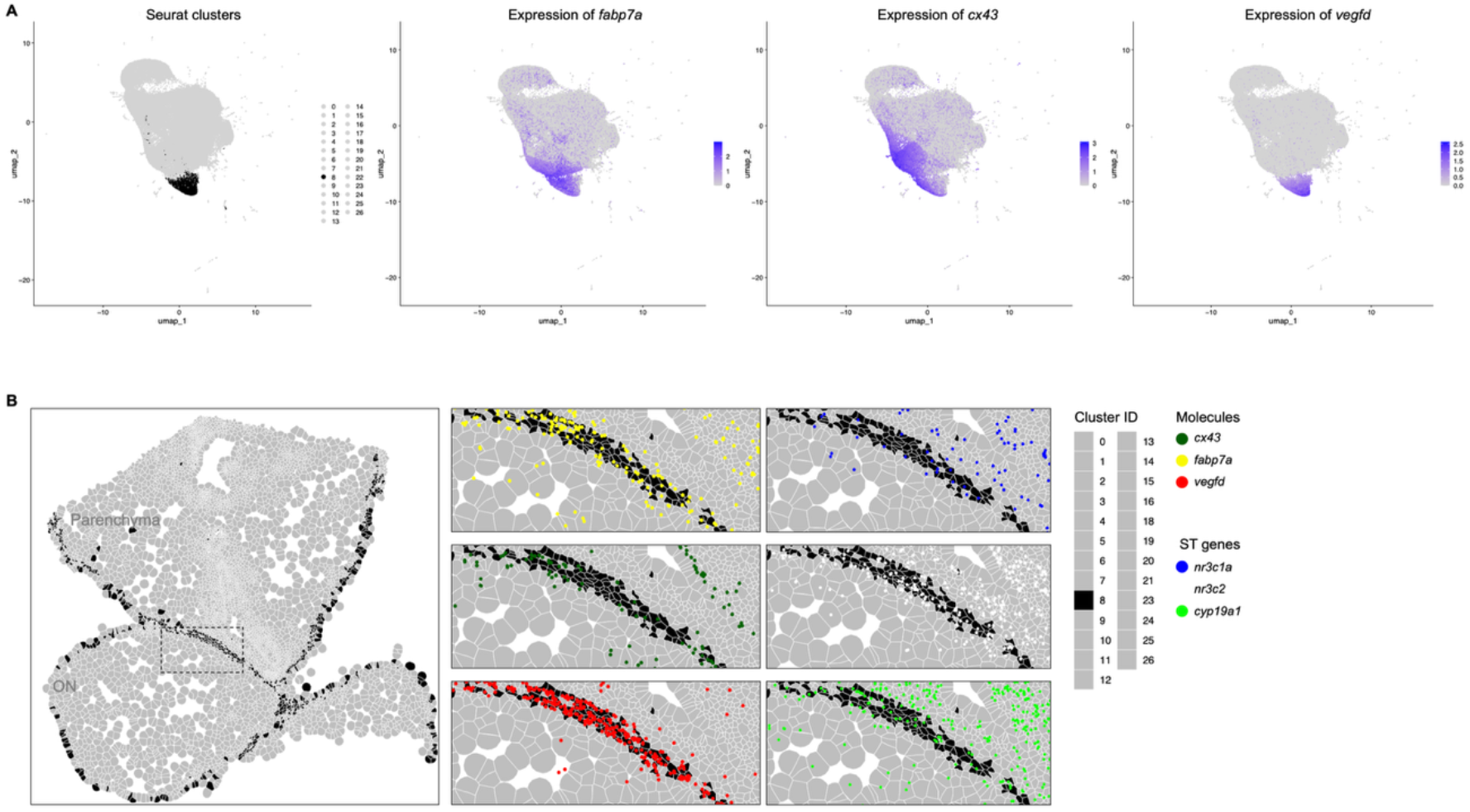
Spatial transcriptomics revealed mLSCs in the *A. burtoni* brain. (A) UMAPs representing M-C8 cells in the male dataset. These cells overexpress *fabp7a, cx43* and *vegfd* marker genes. (B) Cells are located exclusively at the outer boundaries of the parenchyma and the ON (optic nerve). The genetic signature and location of these cells suggest they are mLSCs.

#### Microglia

Several clusters in both sexes corresponded to microglial populations defined primarily by *rps29* together with canonical markers including *vsir, nrros*, and *fgl2a* (Fig. S20). Disease-associated and homeostatic-like microglia were concentrated mainly in posterior hypothalamic nuclei, particularly the NDILI, PGc, CM, and PGl. Some *rps29*+ clusters also co-expressed oligodendrocyte and neuronal markers, suggesting heterogeneous populations containing metabolically active glia and neurons. In both sexes, these microglial populations additionally expressed low levels of *cyp19a1, nr3c2*, and *egr1*, indicating steroid sensitivity and activity-dependent signaling.

#### Neurons

Glutamatergic neurons were spatially restricted to the PGc, habenulae, PGl, and TL, consistent with previous teleost studies *(89, 97)* (Fig. 7A). These neurons strongly expressed *slc17a7a* together with *gabbr2*. Many also co-expressed glutamatergic receptor genes and *gad1*, consistent with hybrid excitatory/inhibitory neuronal states (Fig. 2B) *(13)*. In both sexes, neuron subsets expressed *cyp19a1b, nr3c2*, and *egr1*, revealing that steroid-responsive excitatory neurons express *cyp19a1b*. This contrasts with previous reports localizing *cyp19a1b* to glia in teleosts *(98–101)*.

**Fig. 7.**
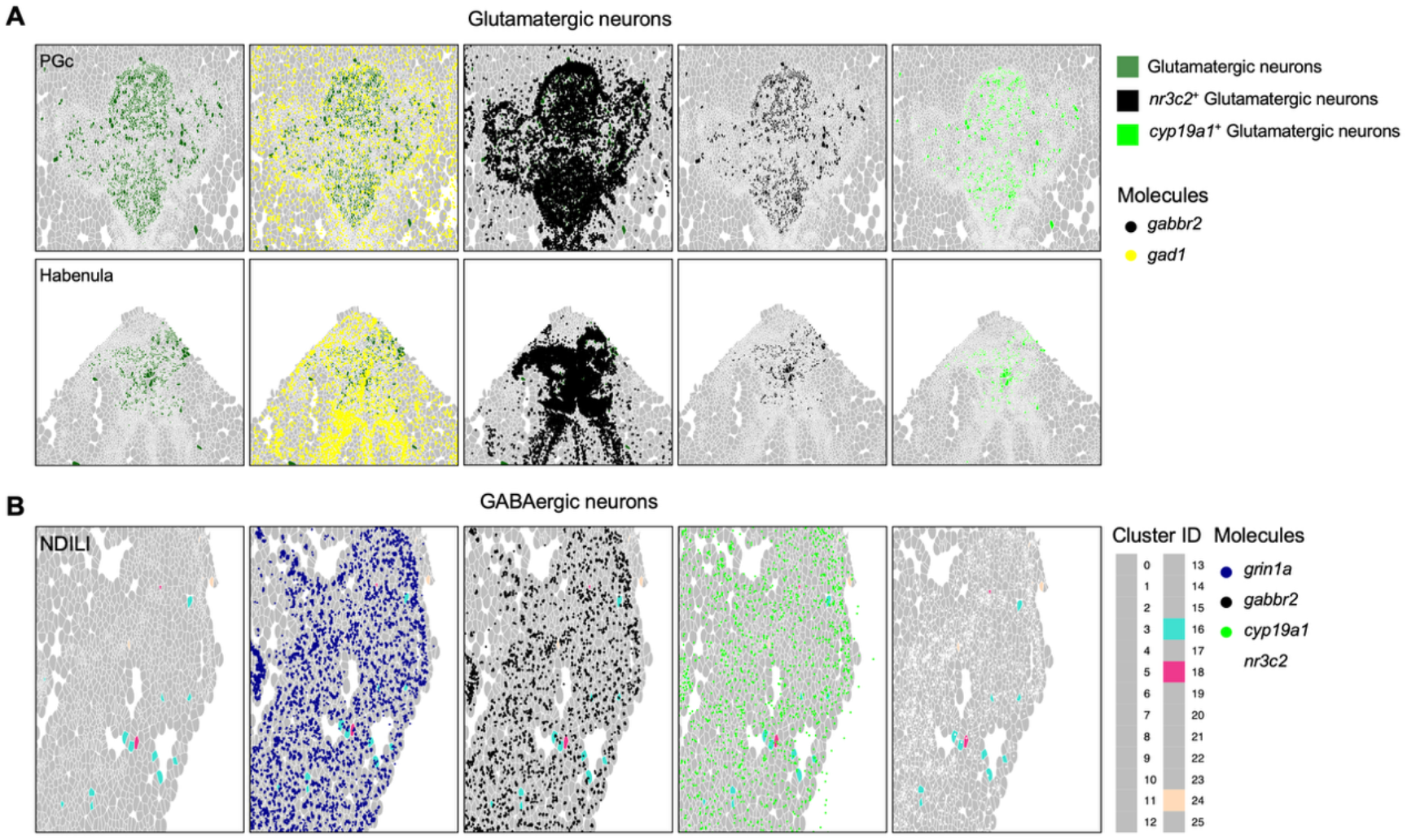
GABAergic and glutamatergic neurons are steroid sensitive and synthesize estradiol. (A) Glutamatergic neurons in the male hypothalamus, here pictured in the PGc and the habenulae. The majority of these neurons express *gabbr2* (>85% of cells). A small proportion of these neurons also express *gad1. Nr3c2* and *cyp19a1* are the two steroid related genes that are the most expressed in glutamatergic neurons (in >50% and >37% of cells, respectively). (B) Only three clusters express *gad1* in 100% of cells, in the female hypothalamus. These GABAergic neurons also express *grin1a* and *gabbr2* (in a maximum of 30% of cells). Steroid related genes are also expressed, predominantly *cyp19a1* and *nr3c2*. See Table S1 for abbreviations.

In contrast, GABAergic neurons were broadly distributed throughout the hypothalamus, especially in posterior nuclei including the NDILI, NDIL, CM, and PGc (Fig. 7B), and similarly expressed *cyp19a1b, nr3c2*, and *egr1* at low levels. Beyond canonical glutamatergic and GABAergic neurons, we identified small neuronal populations enriched for glutamate and/or GABA receptor subunits despite weak *slc17a7a* or *gad1* expression, suggesting neurotransmitter-responsive states concentrated in posterior hypothalamic regions. Large heterogeneous neuronal clusters in both sexes contained mixtures of GABAergic neurons and neurons responsive to both excitatory and inhibitory signaling. These broadly distributed populations also expressed neuropeptide and steroid-related genes, more specifically *cyp19a1b, nr3c2, egr1*, and lower levels of *avp, npy, sst1.1, esr2a*, and *nr3c1a*. The male dataset additionally contained a distinct neuronal population localized almost exclusively to AVT- and GnRH1-rich nuclei (Fig. S21). These cells expressed *gpx3, gad1, rps29* and *stmn2*, consistent with a neuroblast-like identity, together with high levels of *cyp19a1b, nr3c2, avt, gnrh1, sst1.1*, and *egr1*.

F-C7 showed high expression of markers that correspond to diverse cell types *(fgl2a, grin1a, gabbr2, rps29, cacna2d2a, mbpa, etc*.). Subclustering did not clearly resolve distinct cell types in this cluster. Using a PCA-based analysis, we highlighted a transcriptional gradient. As such, cells within hypothalamic nuclei exhibited high PC1 values and were associated with genes such as *grin1a, gabbr2, cacna2d2a, and grin2ab*, suggesting a consistent neuronal identity. On the contrary, cells in the parenchyma, the ON, and those near the outer parenchyma boundaries were characterized by high expression of *rps29, nrros*, and *fgl2a*, suggesting that these cells might be microglia. Another cluster, M-C6, was highly heterogeneous. As in the female, the PCA-based analysis revealed that some of these cells are neurons, located in all nuclei. In contrast, other cells, characterized by high expression of *rps29, vsir*, and *nrros* (i.e., potential microglial cells as well), were localized close to the outer borders of the parenchyma (see Fig. S22).

### Dispersion of ST genes

We mapped the spatial distribution of ST genes to identify hypothalamic populations capable of steroid signaling and local estradiol synthesis (Fig. S23). Androgen receptor genes *ar1* and *ar2* were concentrated primarily in the nPPa, ATn, and ventricular zones, with additional expression in mLSCs and RGC/tanycyte/astrocyte populations (Fig. S24). *Pgr* expression was sparse and localized mainly to the nPPa, and a few transcripts in the NDIL, and NLT. In contrast, *cyp19a1b* was broadly distributed and highly expressed across many neuronal and glial populations, including glutamatergic and GABAergic neurons, GnRH1/AVT neurons, radial glia, and mLSCs, extending well beyond ventricular zones.

*Esr1* expression was sparse and broadly dispersed, whereas *esr2a* and *esr2b* were enriched in the ATn, NLT, NRL, nMMp and nPPa, with *esr2b* particularly enriched in GnRH1/AVT neurons. Glucocorticoid receptor genes also showed distinct distribution. *Nr3c1a* was concentrated in the PGc, NRL, DP, and NLT, whereas *nr3c2* was among the most abundant ST genes, broadly distributed throughout the posterior hypothalamus and strongly enriched in glutamatergic neurons and neurons responsive to glutamatergic and/or GABAergic signaling. The male and female datasets showed largely similar ST gene organization.

### Dispersion of PEP genes and *egr1*

We similarly mapped PEP genes and the activity-dependent marker *egr1* across the hypothalamus (Fig. S25). *Avt*+ cells showed the expected distribution within the nGMp, nMMp, and nPPp. Still, they were also detected in the habenulae, VMn, and E. *Gnrh1* expression was concentrated in the nPPa and in the pineal stalk. In contrast, *npy* and *sst1.1* were enriched primarily in anterior hypothalamic regions, including the Vl, Vd-r, Vv, and Vp, with additional expression in the NLT, VMn, and neighboring nuclei. *Egr1* transcripts were abundant throughout the posterior hypothalamus, particularly in the CM, NRL, NLT, habenulae, NDILI, PGc, and TL. Spatial organization was largely similar between sexes.

### Functional validation of *sst1.1* as a regulator of dominance traits

Next, we aimed to demonstrate the experimental tractability of our *A. burtoni* hypothalamus atlas. Expression of *sst1.1* was higher in DOM GABAergic neurons compared to SUB GABAergic neurons (Fig. 3A). Previous work showed that somatostatin (SST) neurons are larger in DOM compared to SUB *A. burtoni(81)*, and pharmacological manipulation of SST receptors suggests it inhibits aggression. We used CRISPR/Cas9 gene editing to engineer *A. burtoni* lacking functional *sst1.1* (Fig. S26). We propagated an allele with a 70-bp deletion (Fig. 8) and developed a stable breeding line. *Sst1.1* KOs were significantly larger than wild-type (WT) (Fig. 8A), supporting the hypothesis that elevated SST in DOM males shifts energy expenditure away from growth and toward reproductive and territorial defense activities *(81)*. This hypothesis would have been difficult to test pharmacologically because it would require prolonged treatment with unknown doses of agonists or antagonists, substantially reducing experimental throughput. *Sst1.1* KO males were additionally assayed for aggression in a mirror assay *(7, 102)*, a suitable approach for assaying fish that inherently differ in size as a function of the focal independent variable. There were no differences in aggression between KOs and WTs (Fig. 8B).

**Fig. 8.**
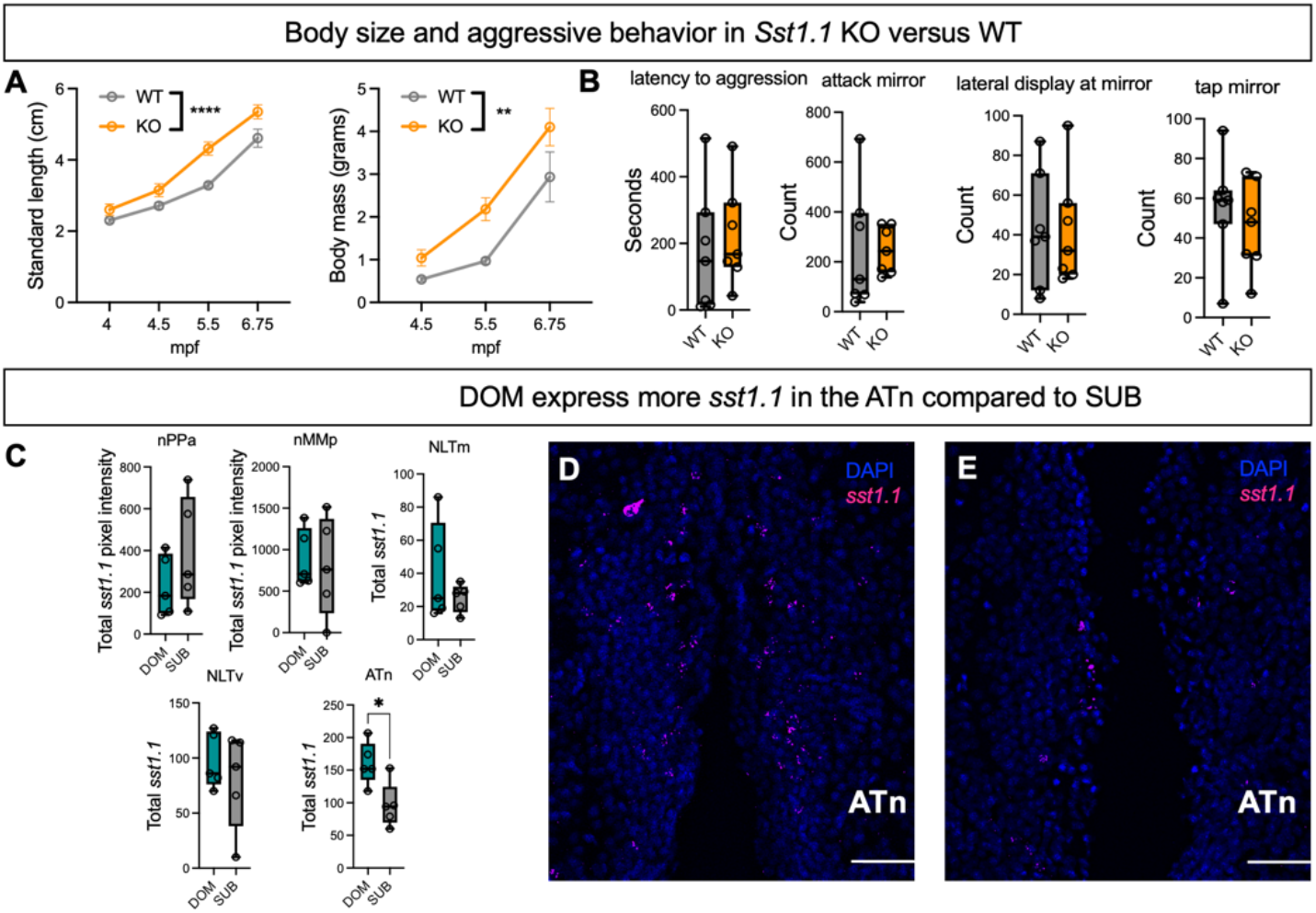
*A. burtoni* hypothalamus-atlas-guided genetic dissection of somatostatin’s role in dominance. (A-B) *sst1.1* KO were larger than WT. (B) There were no differences across multiple aggression variables. (C) DOM males expressed more *sst1.1* than SUB in the ATn but not other hypothalamic regions. (A) Circles plus error bars represent the mean ±SEM at each time point. (B-C) Circles are datapoints for each fish. Bars represent the median with interquartile ranges. (D) Scale bar=50 _μ_M. * p<0.05; ** p< 0.01; **** p<0.0001.

We next aimed to spatially resolve status-dependent differences in *sst1.1* expression in the male hypothalamus. In the spatial dataset, *sst1.1* expression was densest in the NLT and ATn, areas that also showed localized expression of *ar1, esr2a*, and *cyp19a1b* (Fig. S23, Fig. S25). DOM males had higher *sst1.1* expression in the ATn compared to SUB males (Fig. 8). The fish ATn is the homolog of the ventromedial hypothalamus (VMH) in mammals, which regulates metabolism, thermoregulation, and energy expenditure *(103–105)*. Estrogen signaling in the VMH of mice has been shown to modulate metabolic activity and energy expenditure *(106, 107)*, and SST+ neurons in the VMH express estrogen receptors in rats *(108)*. These findings suggest ATn *sst1.1* neurons are a central neuroendocrine node in regulating metabolism and energy allocation in relation to social dominance in *A. burtoni*.

## Discussion

Using scRNA-seq and spatial transcriptomics, we created a spatially resolved cell-type-specific molecular atlas of the *A. burtoni* hypothalamus. Our approach identified major cell classes and subclasses, as well as their topography. Together, these approaches resolved the identities of hypothalamic cells and their spatial organization. The scRNA-seq results identified highly modular, cell-type-specific expression of steroid signaling genes, revealing cell-type-specific sensitivity to social and reproductive state. Spatial transcriptomics enabled us to generate a map of several steroid-signaling and peptide genes in the hypothalamus. Through our *sst1.1* experiments, we showed that our atlas can prospectively guide functional studies. Together, we have established a functionally actionable hypothalamic atlas in *Astatotilapia burtoni*.

The hypothalamus is central to the regulation of social and reproductive behavior across vertebrates. Yet the cellular organization that supports behavioral plasticity, especially in teleost fish, has remained difficult to resolve. By integrating single-cell and spatial transcriptomic approaches in a socially dynamic vertebrate, this study provides a cell-type–resolved framework for understanding the cellular basis of sexually dimorphic physiology and behavior and how social experience and reproductive state reshape hypothalamic state in teleosts. Our findings support a distributed organization in which social, sex-specific, and reproductive signals engage distinct neuronal, glial, and progenitor populations. A key feature of this organization is the modular deployment of endocrine sensitivity. Such an arrangement may allow endocrine signals to differentially influence behavior, physiology, and plasticity.

Social status, sex, and reproductive state engaged overlapping but distinct transcriptional programs across the hypothalamus. Differences between DOM and SUB males included neuropeptide genes and genes associated with neurogenesis, growth, and metabolism, consistent with coordinated changes in hypothalamic cellular state. Reproductive state in females preferentially modulated progenitor and early neuronal populations, linking internal physiological condition to cellular remodeling. Together, these patterns suggest that hypothalamic plasticity arises from distributed changes in cellular state. Persistent neurogenic populations in the adult cichlid hypothalamus further support this view. These progenitor populations exhibited distinct transcriptional responses to social and physiological variables. Spatial transcriptomic analyses anchored these molecular identities within hypothalamic anatomy and identified organizational features shared with other vertebrates despite species-specific differences in gene repertoires and neurogenic capacity.

scRNA-seq and spatial transcriptomics differed in their ability to resolve specific hypothalamic cell populations. We identified six distinct microglial cell types based on the scRNA-seq data, but these types were not faithfully resolved spatially. This likely reflects differences in sequencing depth and cellular composition across samples. Indeed, microglia can change their state depending on local factors in the brain, consistent with the view that microglia exist along dynamic cellular states. Some cell types, such as mLSCs, were identified only in the hypothalamus due to the spatial resolution of cell-type-specific gene expression. The presence of both *ar1* and *ar2* within mLSCs suggests androgen signaling regulates this cell type. Androgens have been shown to regulate angiogenesis and cerebrovascular responses to injury *(109–114)*. These findings suggest that androgen signaling influences mLSC-associated vascular functions.

Guided by transcriptional data, we engineered *sst1.1* mutants using CRISPR/Cas9. As expected, *sst1.1* KOs were larger than WTs. *Sst1.1* expression differences in the ATn between DOM and SUB suggest SST in this region orchestrates metabolic and energy allocation associated with optimal DOM status. We did not observe a difference in aggression between KOs and WTs. In previous pharmacological studies in *A. burtoni*, males injected with the somatostatin antagonist cyclosomatostatin exhibited increased aggression *(115)*. Collectively, these data suggest that distinct SST genes and/or specific SST receptors differentially regulate behavior and energy expenditure in male *A. burtoni*. This experiment illustrates how our atlas can generate testable hypotheses that bridge molecularly defined cells and behavior.

These findings support a distributed cellular model of hypothalamic plasticity in a socially dynamic vertebrate. The distributed and modular organization of hypothalamic plasticity revealed here suggests that social behavior emerges from coordinated functional molecular switches across multiple cell types. Comparative cellular atlases should clarify how conserved hypothalamic circuits generate flexible social behaviors across vertebrates.

## Materials and Methods

### Ethics

All experimental procedures were conducted according to the ethical guidelines for the care and use of laboratory animals. Experiments were done in accordance with The University of Houston Institutional Animal Care and Use Committee (IACUC, Protocol #202000001).

### Animals

Adult *A. burtoni* derived from a wild-caught population from Lake Tanganyika, Africa *(116, 117)* were used in this experiment. Fish were maintained in recirculating systems simulating their natural equatorial environment (25°C, pH = 8.2, photoperiod 12 h:12 h (L:D), constant aeration).

Adult fish were maintained in aquaria with gravel-covered bottoms and terracotta pots cut in half to serve as shelters and spawning territories. Fish were fed *ad libitum* during the day with commercial cichlid pellets (Ken’s Premium Cichlid Pellets 1.5, KensFish, Taunton, MA, USA) and supplemented with brine shrimp (Bio-Pure Frozen Brine Shrimp, Hikari Sales USA, Inc.). Offspring [fry aged between 0 – 12 days post fertilization (dfp)] were collected and placed in a shaking incubator until their yolk sac was resorbed. Then, larvae were transferred into tanks in one of the recirculating systems, where they were fed with powdered pellets supplemented with Freeze-dried cyclops (San Francisco Bay Brand, Inc., Newark, CA) until they were large enough to feed on pellets.

### ScRNAseq

#### Tissue Preparation and Cell Dissociation

Fish were euthanized by immersion in an ice bath followed by cervical transection. Brains were rapidly dissected in ice-cold phosphate-buffered saline (PBS) and transferred to wells containing ice-cold artificial fish cerebrospinal fluid (afCSF) *(118)*. For embedding, 4% ultra–low melting point agarose (Invitrogen) was heated until fully dissolved to yield a clear, homogeneous solution, then cooled on ice with temperature monitored by a thermometer. When the agarose reached ~30°C, a thin layer was poured into a custom mold (Fig. S27), and the brain was positioned on top using a sterile pipette tip to align orientation. A second layer of agarose was applied to secure the tissue, and the sample was reoriented as needed. After solidification, phosphate-buffered saline (PBS) was added to cover the brain, and excess agarose was trimmed away with a chilled razor blade. Regions of interest were microdissected under a stereomicroscope using sterile blades and immediately transferred to 15 mL centrifuge tubes containing 5 mL of ice-cold artificial cerebrospinal fluid (aCSF). The surrounding medium was then transferred to a new tube, leaving just enough to cover the tissue, and kept on ice for subsequent steps.

Tissue was digested in 2 mL of freshly prepared sterile papain solution (250 μL papain (Sigma), 100 μL 1% DNase 1 (Sigma), 200 μL L-cysteine (12 mg/mL, Sigma), in 5 mL of artificial fish cerebrospinal fluid (afCSF)))) at 37°C for 20 min, gently triturated ~10 times using a wide-bore 200 μL pipette tip, and incubated for an additional 20 min. This process was repeated using progressively narrower pipette tips, culminating in a fire-polished Pasteur pipette (~1,000 μm opening), until a homogeneous cell suspension was achieved. The suspension was filtered through a 30 μm MACS® SmartStrainer (Miltenyi Biotec) into a new 15 mL centrifuge tube on ice to remove debris and clumps. A minimal volume of 500 μL was maintained to prevent sample loss (~100–150 μL).

The filtered suspension was supplemented with 2 mL of ice-cold DMEM/F12 medium reserved from earlier steps and processed using the Debris Removal Solution (Miltenyi Biotec). The cell suspension was centrifuged at 300 × g for 10 min at 4°C, and the supernatant was removed. The pellet was resuspended in 1,100 μL of cold PBS, followed by addition of 900 μL of cold Debris Removal Solution. After gentle mixing by pipetting, 4 mL of cold buffer was overlaid dropwise to prevent phase mixing. Samples were centrifuged at 3,000 × g for 10 min at 4°C with full acceleration and brake. The upper phases were discarded, leaving the middle layer containing viable cells. The suspension was diluted to 15 mL with cold buffer, gently inverted three times, and centrifuged again at 1,000 × g for 10 min at 4°C.

The final cell pellet was resuspended in 200 μL of cold, sterile 0.04% bovine serum albumin (BSA; Sigma) in PBS by gentle pipetting until fully dispersed. Cell concentration and viability were determined using a Countess 3 Automated Cell Counter (Thermo Fisher). Target concentration was 700–1,200 cells/μL, with a minimum viability of 75%. Because the 10x Genomics platform typically captures ~60% of loaded cells, total input was adjusted accordingly to ensure desired recovery (e.g., >16,700 cells for 10,000 recovered). Cell suspensions were maintained on ice until loading.

#### 3’ Single Cell RNA-Seq Library Preparation and Sequencing

Cell suspensions were spun at 500 rcf for 5 min, washed and resuspended once in PBS (without calcium and magnesium) with 0.04% BSA and loaded onto a 10X Genomics Chip G, following the *Chromium Next GEM Single Cell 3’ Reagent Kits v3.1 (Dual Index)* protocol. Gene Expression (GEX) sequencing libraries were then generated. Libraries were assessed for quality using a High-Sensitivity D5000 tape strip on an Agilent 4200 TapeStation and quantified with a Qubit Flex Fluorometer (Thermo Fisher Scientific). Libraries were pooled and sequenced on a NextSeq 2000 (Illumina).

#### Post-Sequencing Processing

The 10x Genomics CellRanger count pipeline was run on a high-performance computing cluster using CellRanger v7.0.0 across all FASTQ files. The sequencing data was aligned to the *Haplochromis burtoni* AstBur reference genome.

### Analysis of Single Cell Gene Expression Data

Single-cell RNA sequencing GEX data were obtained from four female samples (two gravid, two non-gravid based on visual assessment) and three male samples (two DOM, one SUB based on visual assessment) samples. Filtered matrix files from CellRanger v7.0.0 were used as input into R and converted into Seurat objects using the CreateSeuratObject function from the Seurat package24,25. Seurat objects were created and processed independently for each sample. Each Seurat object underwent SCT normalization using the SCTransform function. The cells were filtered by feature count, retaining those with 500 to 20,000 features. Principal Component Analysis (PCA) and UMAP dimensionality reduction were performed on each object using RunPCA and RunUMAP functions, respectively. FindNeighbors and FindClusters functions were used to define clusters in the data. The DoubletFinder (RRID:SCR_018771) package was deployed on each Seurat object to identify and remove doublets, followed by subsetting to retain only the singlet cells26. The objects were then annotated with the relevant metadata. Following individual processing, the objects were integrated into a single Seurat object using the Seurat package’s data-integration features.

### Annotation of Cell Types

Cell types were assigned based on the expression of canonical marker genes as defined in previous studies and described in detail in the main text.

### Spatial transcriptomics

#### Protocol

We designed a custom 50-gene panel that includes putative cell-type marker genes to spatially resolve the different hypothalamic cell types and confirm marker gene co-expression identified by the scRNA-seq experiment. 35 marker genes were selected based on several criteria. First, we preselected genes according to: (1) the cluster(s) in which they were expressed (ensuring that one or more genes represented all clusters), (2) the proportion of cells expressing each gene within a given cluster, (3) and their expression level in each cluster. We then refined this list by removing genes that were redundant with others and/or had a high risk of optical crowding. We added nine steroid-related (SR) genes and six peptide (PEP) genes, given their relevance to regulating socio-sexual behaviors in vertebrates and to spatially validating their cell-type-specific expression. All genes are described in Supplemental table 1.

Two Xenium slides were processed: one containing 8 brain sections of the socially ascending male (referred to as “M”, GSI = 0.82) and the other slide containing 8 brain sections of a female with moderate gravidity (F1, referred to as “F” in the rest of the manuscript, GSI = 6.79).

Since brains used in this experiment were fixed and frozen, the first steps of the protocol were done following an adaptation from the “Visium CytAssist Spatial Gene Expression for Fixed Frozen – Rehydration (CG000662)” and “decrosslinking (CG000580)”. For the rehydration step, slides were incubated on a thermocycler at 37°C (1 min) and immersed in the following solutions: 1x PBS (5 min), RNAse-free water (3 min), 100% ethanol (3 min), 70% ethanol (3 min), RNAse-free water (20 sec). Slides were then mounted in the Xenium cassette and PBS-T was added onto them. For the decrosslinking protocol, PBS-T was removed from the slides and the decrosslinking buffer was added (prepared following the manufacturer’s instructions, using the “Xenium slides and sample prep kit [PN-1000460], 10x genomics). Slides were incubated on a thermocycler at 80°C (30 min) followed by a re-equilibration step at 22°C (10 min). The decrosslinking buffer was then retrieved and the slides were rinsed with PBS-T (2 × 1 min).

Subsequently, slides were immediately processed according to the “Xenium In Situ Gene Expression - Probe Hybridization, Ligation & Amplification (CG000582, Rev E)” workflow. This protocol included a few steps briefly described here. First, slides were incubated on a thermocycler at 50°C (24 hours) with our 50-gene custom panel, such that the probes could hybridize to the tissue. This was followed by a post hybridization wash (37°C, 30 min). After unbound probes were removed, the junction between the hybridized probe ends was sealed using a ligase (37°C, 2 hours). Once this ligation step was done, probes were enzymatically amplified (30°C, 2 hours). Finally, slides were washed and processed for autofluorescence quenching and nuclei staining. Reagents used for these steps all belong to the “Xenium slides and sample prep kit (PN-1000460)” from 10x Genomics.

Then, the Xenium instrument was loaded with the following: Xenium decoding reagent module A (1000624) and module B (1000625), all Xenium sample wash buffers (3001198, 3001199, 3001200), the Xenium probe removal buffer (3001201), the objective wetting consumable, and the extraction top and the pipette tip rack (all 10x genomics). Information regarding the slide was added to the instrument (e.g., slide ID, JSON file, etc.). Both slides were loaded into the instrument. A sample scan was then performed, allowing to select the Regions of Interest (ROI) on the sections. We selected 16 ROIs in total (i.e., 8 on each slide), each including an entire brain section (containing between 15 and 49 Fields Of View [FOVs]).

After completion of the run (about 35 hours), output information was retrieved from the instrument. Output files include (1) an analysis summary of the data (for each ROI) and (2) an image of each ROI in which nuclei where transcripts and cell clusters could be seen using the Xenium Explorer software (version 3.1.1, 10x genomics) with default cell segmentation. All data were stored in a hard drive for further analysis. Slides were carefully removed from the Xenium instrument, covered with PBS-T, and stored in the dark at 4°C until H&E staining (Vector Labs, H-3502)

#### Data analysis

All eight tissue sections spanning the hypothalamus were processed and analyzed for each individual in RStudio (version 2025.09.0+387, Cucumberleaf sunflower) using Seurat (v5.20.1). As previously described, entire brain sections were processed on the Xenium instrument. Since our analysis focused solely on the hypothalamus, we sought to specifically subset this region. To do so, we manually drew hypothalamic boundaries in Xenium Explorer and imported these coordinates into R using a custom function for spatial subsetting, resulting in eight cropped sections including the hypothalamus only in each fish.

Cells with 0 counts and transcripts with a QV less than 20 were removed from the dataset. SR and PEP genes were temporarily excluded from the dataset to ensure they did not influence dimensionality reduction or clustering, which are solely based on the 35 marker genes selected from the scRNA-seq results. Cropped sections were merged into a single data object, allowing all eight hypothalami to be processed and analyzed as a whole.

Before dimensionality reduction, we conducted preliminary analyses to determine the optimal number of principal components (PCs) that capture at least 90% of the variance in the male and female datasets. These analyses indicated that 16 PCs were sufficient for the male dataset, and 18 PCs for the female dataset. All subsequent clustering analyses were performed using a fixed resolution parameter of 0.3.

Then, data were normalized using SCTransform, followed by dimensionality reduction with a principal components analysis (PCA) and visualization with uniform manifold approximation and projection (UMAP). Cell clustering was performed using shared nearest-neighbor graphs, and clusters were visualized with a custom color palette to ensure consistency across all figures. Average gene expression per cluster was calculated to summarize transcriptional profiles across clusters. These expression values were scaled by gene and visualized as a heatmap, to highlight differences in relative expression patterns among clusters.

To facilitate quantitative and spatial analyses, several custom R functions were developed to extract and visualize expression data from our dataset. These functions were used to calculate, for each hypothalamic section, the total number of cells, the number of cells belonging to a given cluster, the number and percentage of cells expressing a target gene, and the proportion of cells co-expressing genes of interest. Additional functions were implemented to visualize RNA molecules directly on tissue sections, either alone, alongside other transcripts, or overlaid on specific cluster cells. Custom plotting functions were also created to highlight cells co-expressing at least 2 genes, allowing the spatial distribution of gene co-expression to be mapped in each hypothalamic section.

After completing clustering using the 35 marker genes, the SR and PEP genes could be reintroduced into the dataset (i.e., in the merged dataset of all 8 hypothalami) for subsequent visualization on tissue sections to examine their spatial distribution in the hypothalamus.

#### Sub-clustering

Clusters that appeared heterogeneous (highly dispersed on the UMAP and contained several highly expressed marker genes) were reanalyzed at higher resolution using subclustering. For each parent cluster, cells were subsetted. Preprocessing was comprised of log-normalization, scaling, and dimensionality reduction (PCA) using the optimal number of PCs determined for that parent cluster. Finally, a UMAP and a dot plot (using log-normalized RNA expression data) were generated for visualization.

#### Spatial heterogeneity within clusters

To characterize spatial heterogeneity of gene expression within individual clusters, we performed a cluster-specific PCA. Scores for the first principal component (PC1) were assigned to individual cells and visualized on the brain sections by coloring cells according to their PC1 values. Genes contributing to the observed variation were identified based on their loadings on PC1.

### Generation of *sst1.1* mutants using CRISPR/Cas9 gene editing

Mutations were introduced into a coding region of *sst1.1* using CRISPR following previously published protocols *(119, 120)*. Briefly, two crRNA target sites were selected from top candidates in CHOPCHOP that recognized sequences 5′ of predicted essential domains and were synthesized by IDT (Fig. S26). tracrRNA was synthesized by IDT (1072532). The crRNA:tracrRNA duplex (final conc. 5 μM) was delivered along with Cas9 protein (IDT, final conc. 5 μM) into single-cell embryos within 90 minutes of fertilization. Mosaic F_0_ fish were screened for extent of mutation in finclip genomic DNA, and fish with high mutation rates and large indels as shown by fragment analysis were selected for out-crossing to unrelated WT fish to isolate large, frameshift indels. A subset of heterozygous offspring showed a 70-bp deletion, which we propagated further to unrelated WT fish. Heterozygous fish were then in-crossed to generate WT, heterozygous, and homozygous null (KO) fish. A subset of crispants and their uninjected siblings were subjected to a mirror assay (see below).

### Quantification of behavioral and physiological phenotypes in *sst1.1* crispants and mutants

Seven *sst1.1* KO were assayed for aggression in a mirror assay as we have done previously *(7, 102)*. Briefly, the mirror assay consists of a white tub (Sterilite; 400 mm × 318 mm × 12 mm) filled 6 cm deep with UV-sterilized aquaria water with a mirror replacing one side of the assay tub. An opaque cover was first placed on the mirror for a 15-min habituation period to allow subjects to acclimate to the assay tank. The opaque cover was then removed, and 30 min of behavior was recorded. We also assessed body-size traits in *sst1.1* KOs and WTs. Fish were netted and then patted dry before measuring standard length (SL) and body mass (BM).

### Measuring the *sst1.1* expression in the hypothalamus of DOM versus SUB males

We performed HCR to detect *sst1.1* expression in brain tissue from DOM and SUB males. To establish DOM males, we housed them in a stable DOM tank paradigm, which reliably induces dominance-related behavioral and physiological traits *(121, 122)*. Briefly, five size-matched males (n=5 DOM) are housed in a 121-liter tank with 10 to 15 females and five potential mating sites that are represented by halved terra cotta pots. To establish SUB males, we used a social suppression paradigm, which reliably induces the full suite of SUB traits. To create SUB fish, five males (n=5 SUB) were housed with three larger DOM suppressor males and 10-15 females.

After four to six weeks in their status-inducing tanks, focal fish were removed, euthanized by immersion in an ice-water bath, weighed (body mass, BM), and measured for standard length (SL). After cervical transection, brains were exposed from the olfactory bulbs to the spinal cord and fixed in the head in 4% PFA (pH 7.4) for 24 h at 4°C. Heads were then rinsed in 1× PBS (3 × 5 min, then ~6 h at 4°C) and cryoprotected in 30% sucrose overnight at 4°C. Brains were extracted from the heads, embedded in Neg-50 (22-046-511, Fisher Scientific) and kept at −80°C until sectioning. Gonads were removed from the visceral cavity and weighed (gonads mass, GM) to calculate the gonadosomatic index [GSI = (GM/BM) × 100]. Brains were sectioned in the coronal plane using a cryostat (Thermo Scientific, HM 525 NX) at a thickness of 20 μm on three alternate slide series. Slides were air-dried for 24 h then stored at −80°C until further processing.

HCR was performed according to manufacturer protocols, recommendations, and reagents, unless otherwise noted. In summary, on day 1 of the protocol, tissue slides were exposed to LED light for 60 min to reduce autofluorescence. Subsequently, tissue was immersed in 0.2 % Triton X-100 (in 1× PBS) (Thermo Fisher; Gibco) for 30 min to increase signal detection. Next, tissue was dehydrated with serial concentrations of EtOH (Thermo Fisher) at 50 %, 70 %, and 100 %, before adding Proteinase K (1:2000) (Thermo Fisher) in 1×PBS. HCR probes were then hybridized to tissue by adding 16nMs of RNA probe in hybridization buffer for 12-16 h at 37 °C. We used the HCR probes against *sst1.1* and *grin2ab* (for a separate experiment) that were engineered by Molecular Instruments. Following the hybridization step, specific probe amplifiers tagged with distinct fluorophores were added at 60 nM into a single amplifier solution. Slides were incubated in amplifier solution at room temperature in a dark chamber overnight. A 647 amplifier was used to detect the *sst1.1* probe, while a 488 amplifier was used to detect the *grin2ab* probe. Slides were then washed in 5× SSCT (0.1%Tween-20) (Gibco; Thermo Fisher), and cover slipped using Prolong Gold mounting media with DAPI (Invitrogen). Final sample sizes for cell quantification: WT: n = 5; Mut: n = 5; and Empty: n = 5.

Slides were imaged using a Nikon Eclipse 80i Microscope MicroFire™ at 20× magnification in cy5 and DAPI wavelength filters in the using multichannel, multiplane image capture. Our goal was to quantify the amount of *sst1.1* in the hypothalamus. *Sst1.1* expression was observed in the NLTv, NLTm, ATn, and POA. *Sst1.1* puncta could be clearly delineated from one another in DAPI+ cells in all regions except the POA, where expression was too intense for individual counting of puncta. To count puncta in the NLTv, NLTm, and ATn, we used Fiji ImageJ cell-counting software to tally total puncta across all cells. In the POA, cy5 pixel intensity was used to quantify *sst1.1* expression. We quantified *sst1.1* in two subsequent sections per brain region.

### Statistics

DEG analysis was done in Seurat. T-tests were used to compare phenotypic differences between *sst1.1* KOs and WTs with a significance threshold of *p<0.05*.

## Acknowledgments

We thank Lillian Jackson and Mariana Lopez for assisting with dissections in support of the scRNA-seq protocol. We thank Sofia Young for assistance in generating the *sst1.1* mutants.

## Authors contributions

M.C., A.P.H., and B.A.A. designed the scRNA-seq experiment. A.P.H. optimized the scRNA-seq protocol. M.C. and B.A.A. analyzed and interpreted the scRNA-seq data. M.D. and B.A.A. designed the spatial transcriptomics experiment. M.D. and M.C. wrote custom code for analyzing spatial transcriptomics data. M.D. analyzed the spatial transcriptomics data, and B.A.A and M.D. interpreting spatial transcriptomics findings. L.A.S., A.P.H., and B.A.A. engineered the *sst1.1* mutants. LA.S. and B.A.A. designed the *sst1.1* experiments, and LA.S. performed all *sst1.1* mutant experiments and analyzed the data. P.H.G. provided resources through the UH Sequencing and Editing Core. M.D., M.C., and B.A.A. wrote and edited the initial draft of the manuscript. P.H.G. reviewed the manuscript draft. M.D., M.C., and B.A.A. finalized the manuscript. B.A.A. provided the funding that supported all experiments.

## Data and materials availability

FASTQ sequencing files are available on NCBI SRA: Bioproject PRJNA1477979 (to be released upon acceptance for publication). Cluster gene marker data, DEG and GO results tables, and custom code for spatial analysis are available on Dryad (DOI: 10.5061/dryad.7h44j1097). Tables S1-S2 and Figures S1-S28 are in the Supplementary Material

