## Supplementary Material for "Cell-type-specific architecture of the hypothalamus in a socially plastic vertebrate"

**Supplementary Tables S1 and S2**

**Supplementary Figures S1-S27**

| Gene name | Ensemble ID | General class |
|---|---|---|
| *adamts15* | ENSHBUG00000019193 | Astrocytes |
| *aif1* | ENSHBUG00000004663 | Microglia |
| *bcl11ab* | ENSHBUG00000020505 | Neurogenesis lineage |
| *cacna2d2a* | ENSHBUG00000002586 | Neurons |
| *cx43* | ENSHBUG00000001315 | Astrocytes |
| *edaradd* | ENSHBUG00000014521 | Microglia |
| *fabp7a* | ENSHBUG00000006848 | Astrocytes |
| *flg2a* | ENSHBUG00000021176 | Microglia |
| *fybb* | ENSHBUG00000015604 | Macrophages |
| *gabbr2* | ENSHBUG00000019409 | Neurons |
| *gad1* | ENSHBUG00000004335 | Neurogenesis lineage |
| *gem* | ENSHBUG00000016008 | Neurogenesis lineage |
| *gfap* | ENSHBUG00000008793 | Astrocytes |
| *gpx3* | ENSHBUG00000004581 | Neurogenesis lineage |
| *grin1a* | ENSHBUG00000005579 | Neurons |
| *grin2a* | ENSHBUG00000022380 | Neurons |
| *grin2ab* | ENSHBUG00000018585 | Neurons |
| *hbe1* | ENSHBUG00000004147 | Macrophages |
| *mbpa* | ENSHBUG00000018071 | Oligodendrocyte lineage |
| *myrf* | ENSHBUG00000018627 | Oligodendrocyte lineage |
| *nes* | ENSHBUG00000017503 | Neurogenesis lineage |
| *nrros* | ENSHBUG00000015577 | Microglia |
| *p2ry12* | ENSHBUG00000023756 | Microglia |
| *pip5kl1* | ENSHBUG00000013098 | Oligodendrocyte lineage |
| *plekha7* | ENSHBUG00000013514 | Neurogenesis lineage |
| *rps29* | ENSHBUG00000018115 | Microglia |
| *sgsm2* | ENSHBUG00000007037 | Macrophages |
| *slc17a7a* | ENSHBUG00000019189 | Neurons |
| *stim1a* | ENSHBUG00000001129 | Oligodendrocyte lineage |
| *stmn2* | ENSHBUG00000000521 | Neurogenesis lineage |
| *th2* | ENSHBUG00000002626 | Neurons |
| *traf4a* | ENSHBUG00000008510 | Oligodendrocyte lineage |
| *vegfd* | ENSHBUG00000016210 | Tanycytes |
| *vsir* | ENSHBUG00000010891 | Microglia |
| *zeb1l* | ENSHBUG00000009297 | Microglia |
| *ar1* | ENSHBUG00000002755 | ST gene |
| *ar2* | ENSHBUG00000021091 | ST gene |
| *nr3c1a* | ENSHBUG00000022339 | ST gene |

| | | |
|---|---|---|
| *nr3c2* | ENSHBUG00000008760 | ST gene |
| *pgr* | ENSHBUG00000001421 | ST gene |
| *cyp19a1* | ENSHBUG00000011259 | ST gene |
| *esr1* | ENSHBUG00000008433 | ST gene |
| *esr2a* | ENSHBUG00000011554 | ST gene |
| *esr2b* | ENSHBUG00000012640 | ST gene |
| *gnrh1* | ENSHBUG00000013256 | PEP gene |
| *kiss1rb* | NW_024582415.1 | PEP gene |
| *npy* | ENSHBUG00000017245 | PEP gene |
| *sst1.1* | ENSHBUG00000001129 | PEP gene |
| *avp* | ENSHBUG00000003494 | PEP gene |
| *egr1* | ENSHBUG00000002344 | Immediate early gene |

**Table S1.** List of all marker genes used in this study; 35 marker genes selected based on the scRNA-seq results; steroid-related genes (SR genes); and genes encoding peptides (ST genes) and an immediate early gene.

| ***List of abbreviations*** | |
|---|---|
| ATn | Anterior tuberal nucleus |
| CM | Corpus mammillare |
| DP | Dorsal posterior thalamic nucleus |
| Ha | Habenula |
| NDIL | Diffuse nucleus of the inferior lobe |
| NDILI | Lateral part of NDIL |
| NLT | Lateral tuberal nucleus |
| nGMp | Magnocellular preoptic nucleus, gigantocellular division |
| nMMp | Magnocellular preoptic nucleus, magnocellular division |
| nPPa | parvocellular preoptic nucleus, anterior part |
| nPPp | parvocellular preoptic nucleus, posterior part |
| NRL | Nucleus of the lateral recess |
| ON | Optic nerve |
| PGc | Commissural preglomerular nucleus |
| PGl | Lateral preglomerular nucleus |
| TL | Torus longitudinalis |
| Vd-r | dorsal part of the ventral telencephalon, rostral subdivision |
| Vl | Lateral nucleus of the ventral telencephalon |
| VMn | Ventromedial thalamic nucleus |
| Vp | Postcommissural nucleus of the ventral telencephalon |
| Vv | Ventral nucleus of the ventral telencephalon |

**Table S2**. List of abbreviations.

### Supplementary Figures

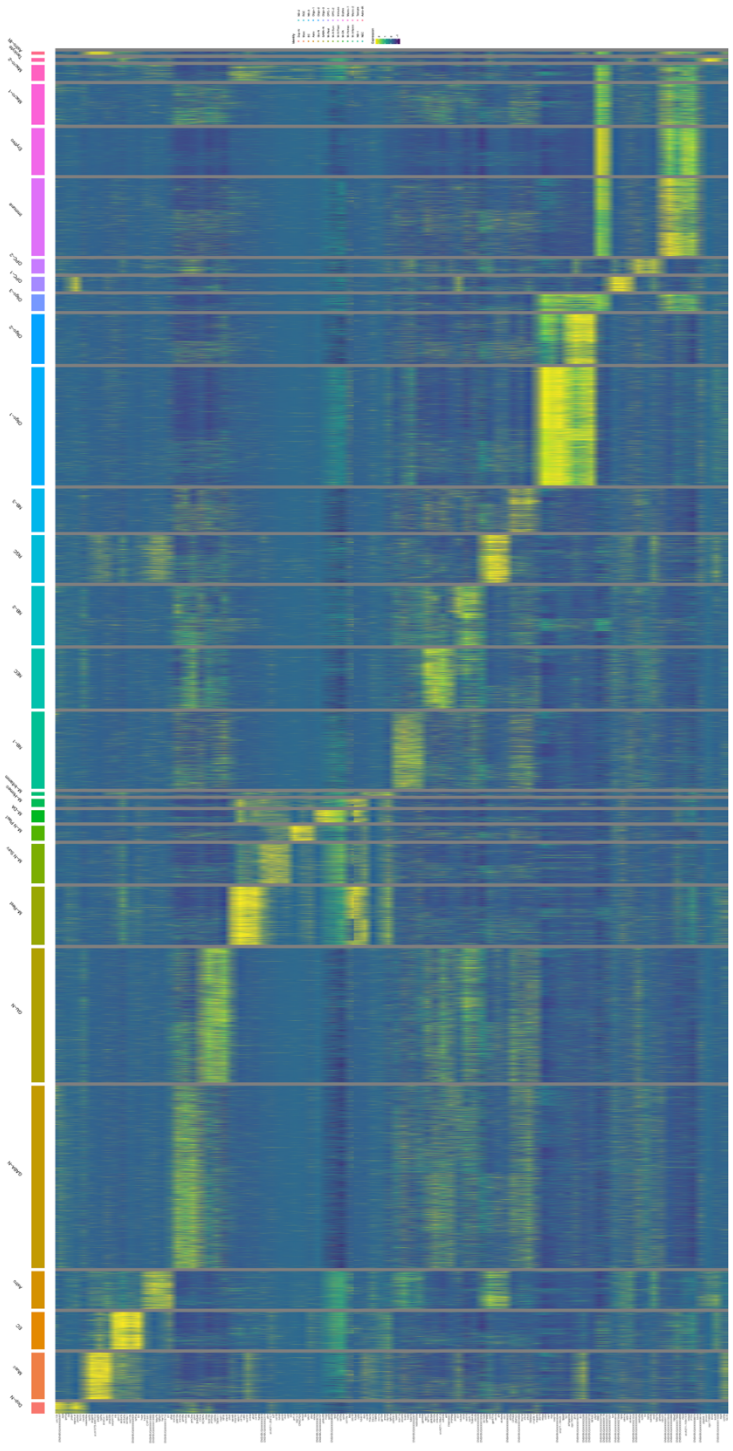

**Figure S1. Heat map showing gene expression across 28 cell type clusters.** The top 10 genes from each cluster were plotted on a heatmap to show the distinct gene programs present. Genes are along the x-axis. Each line is a cell. Bars overlapping the cells demarcate a particular cluster based on gene expression. The color code each cell type and match that present in the UMAP (Fig. 2A). Several genes are written as their Ensemble code given lagging annotation of the *A. burtoni* genome. Ensemble genes can be decoded at ensemble.org. This heat map is also available as a PDF at this link: *https://ucla.box.com/s/eh9azmovdsavicogi9jv75qpp5muplac*

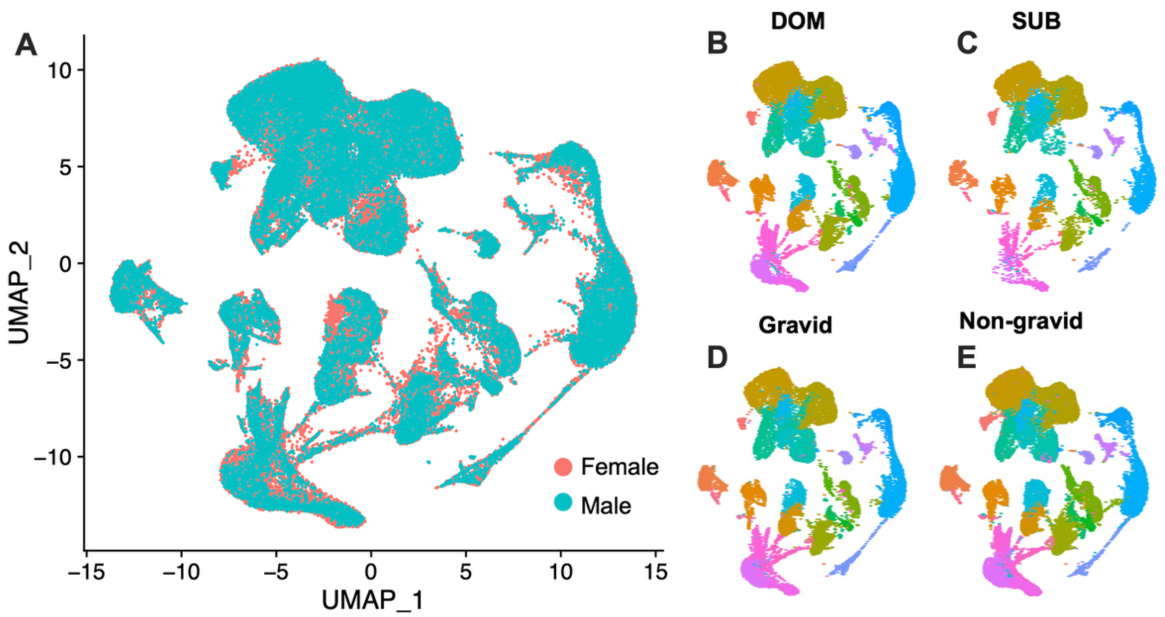


**Figure S2. Uniform manifold approximation and projections (UMAPs) of *A. burtoni* scRNA-seq of the hypothalamus based on sex, status, and reproductive state.** UMAPs demonstrate similar contribution of cells from males and females regardless of status (DOM or SUB) or reproductive state (Gravid or Non-gravid).

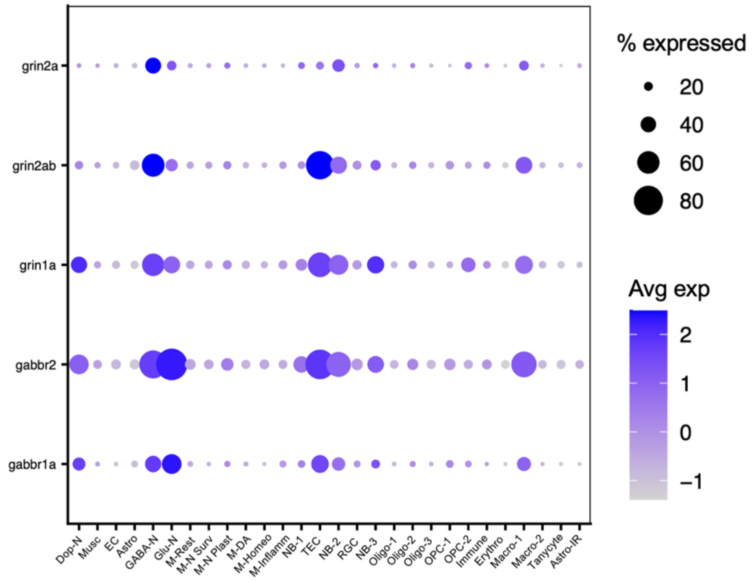


**Figure S3. GABAergic and glutamatergic postsynaptic receptors across neurons, TECs, and neurogenic cells.** GABA and glutamate receptor subunits are distributed across hypothalamic cell types in *A. burtoni*. The dot plot shows genes on the y-axis and the corresponding cell types on the x-axis. N=Neuron; GABA=GABAergic; Glu=Glutamatergic; Dop=Dopamine; NB=Neuroblast; TEC=Tanycyte-like ependymal cells; Oligo=Oligodendrocytes; OPC=Oligo precursor cells; Musc=Muscle; EC=Endothelial cells; RGC=Radial glia cells; Astro=Astrocytes; M=Microglia; Plast=Plasticity; Surv=Survival; Homeo=Homeostatic; DA=Disease associated; Inflam=Inflammatory; Rest=Resting; Macro=Macrophages; Erythro=Erythrocytes.

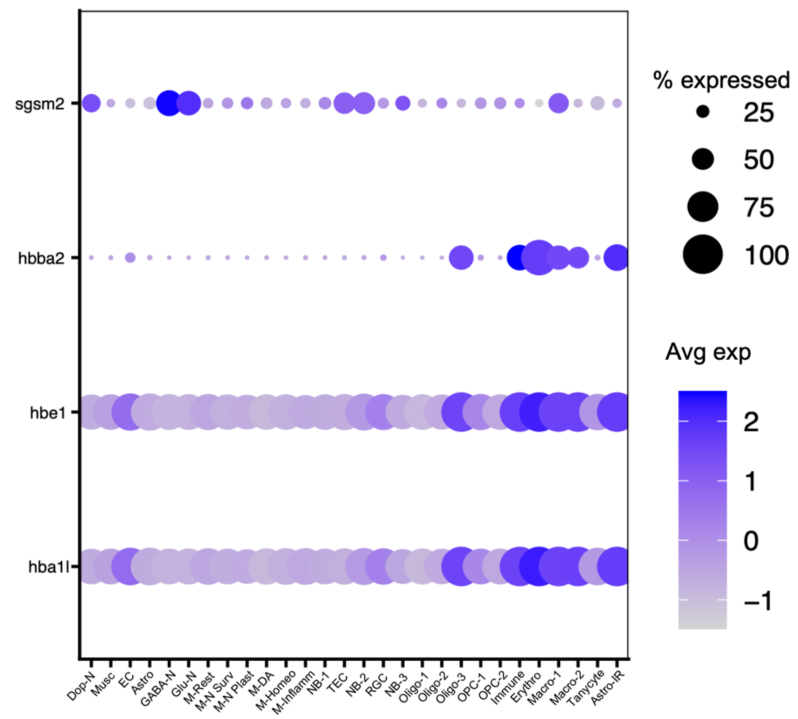


**Figure S4. Specific gene programs define different immune cells.** Macrophages, erythrocytes, and other immune cells are defined by distinct expression patterns. The dot plot shows genes on the y-axis and the corresponding cell types on the x-axis. N=Neuron; GABA=GABAergic; Glu=Glutamatergic; Dop=Dopamine; NB=Neuroblast; TEC=Tanycyte-like ependymal cells; Oligo=Oligodendrocytes; OPC=Oligo precursor cells; Musc=Muscle; EC=Endothelial cells; RGC=Radial glia cells; Astro=Astrocytes; M=microglia; Plast=Plasticity; Surv=Survival; Homeo=Homeostatic; DA=Disease associated; Inflam=Inflammatory; Rest=Resting; Macro=Macrophages; Erythro=Erythrocytes.

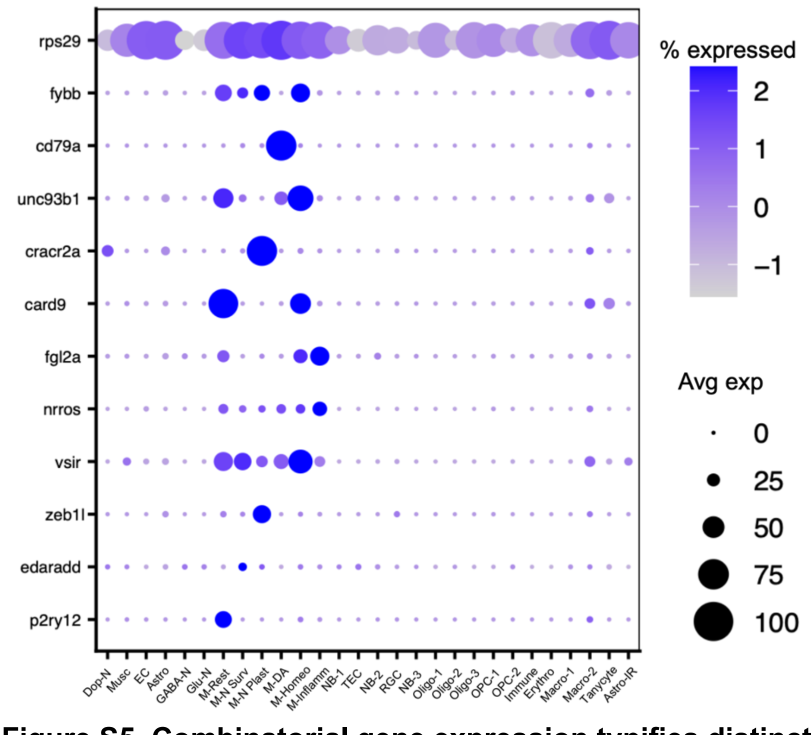


**Figure S5. Combinatorial gene expression typifies distinct microglia states.** Six different microglia types are defined by distinct gene expression patterns. Each gene but *rps29* is distinctly expressed in one or more microglia only. Although *rps29* is present in all cell types, it is relatively elevated in disease associated microglia. The dot plot shows genes on the y-axis and the corresponding cell types on the x-axis. N=Neuron;

GABA=GABAergic; Glu=Glutamatergic; Dop=Dopamine; NB=Neuroblast; TEC=Tanycyte-like ependymal cells; Oligo=Oligodendrocytes; OPC=Oligo precursor cells; Musc=Muscle; EC=Endothelial cells; RGC=Radial glia cells; Astro=Astrocytes; M=Microglia; Plast=Plasticity; Surv=Survival; Homeo=Homeostatic; DA=Disease associated; Inflam=Inflammatory; Rest=Resting; Macro=Macrophages; Erythro=Erythrocytes.

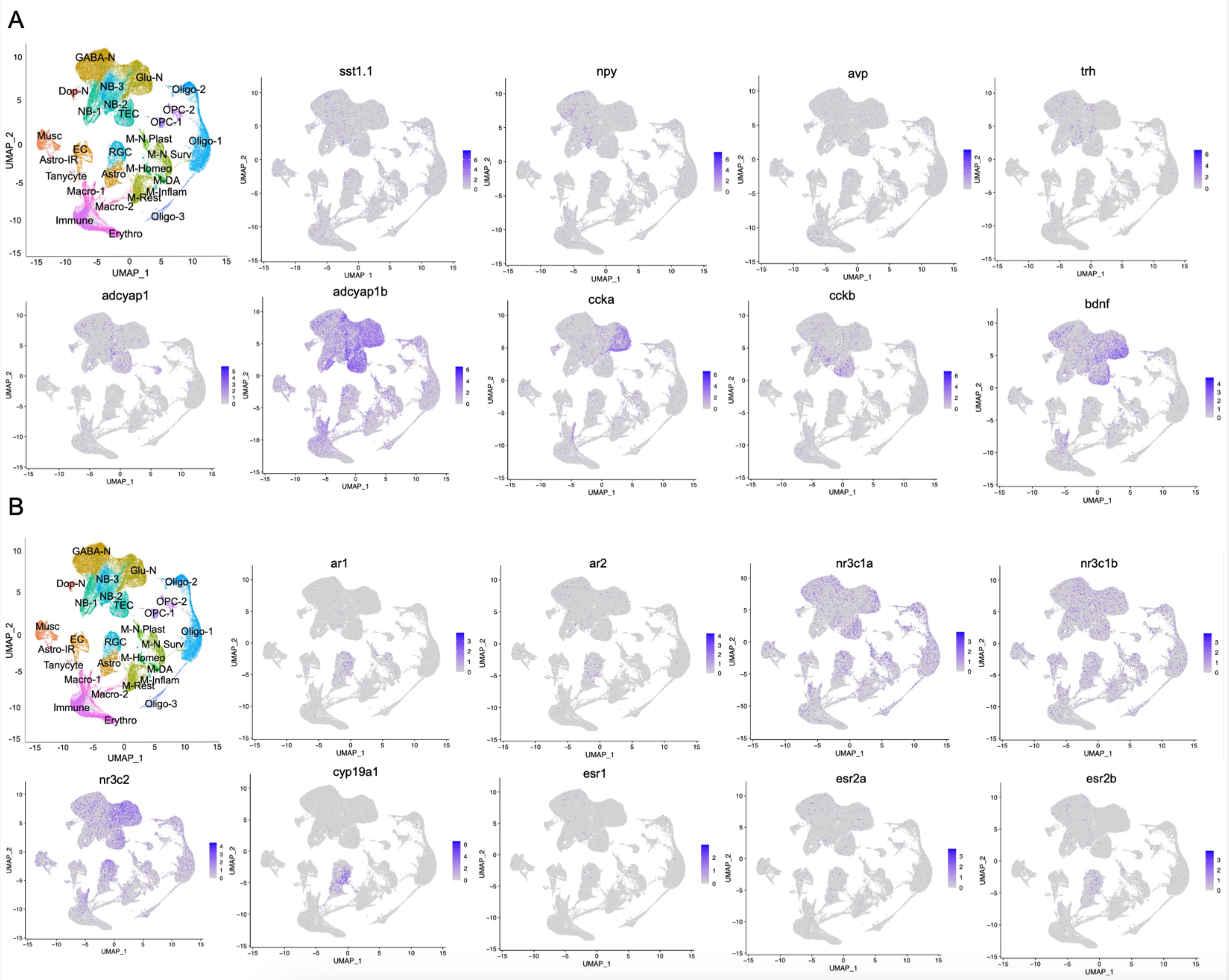


**Figure S6. Peptide and steroid signaling molecule gene expression across cell types.** Feature plots show expression of (A) peptides and (B) steroid signaling genes across cell types. N=Neuron; GABA=GABAergic; Glu=Glutamatergic; Dop=Dopamine; NB=Neuroblast; TEC=Tanycyte-like ependymal cells; Oligo=Oligodendrocytes; OPC=Oligo precursor cells; Musc=Muscle; EC=Endothelial cells; RGC=Radial glia cells; Astro=Astrocytes; M=Microglia; Plast=Plasticity; Surv=Survival; Homeo=Homeostatic; DA=Disease associated; Inflam=Inflammatory; Rest=Resting; Macro=Macrophages; Erythro=Erythrocytes.

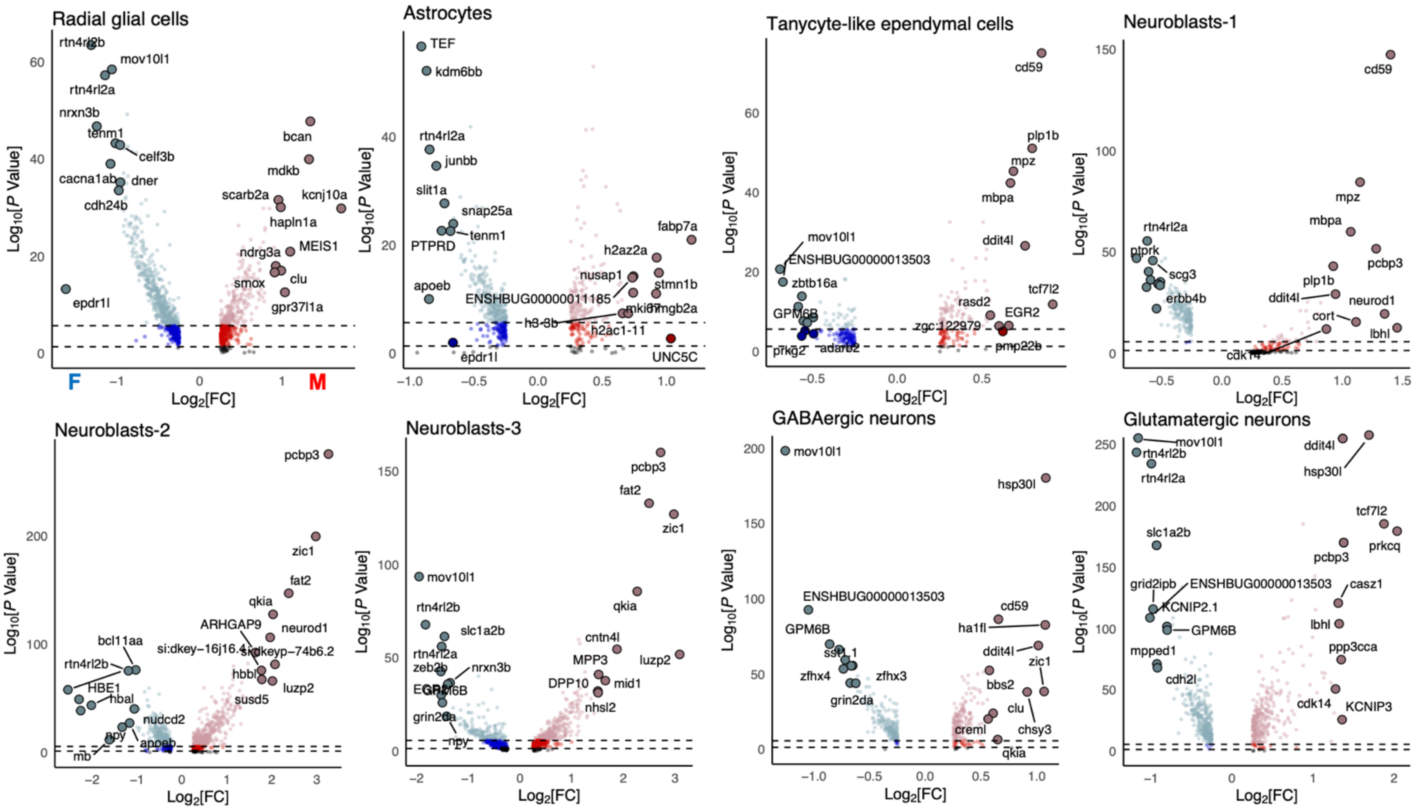


**Figure S7. Sex-dependent cell-type-specific gene expression differences.** Volcano plots of differentially expressed genes (DEGs) as a function of sex. Genes expressed higher in males (M, red font) have positive $Log_2$[FC] values; genes expressed higher in females (F, blue font) have negative $Log_2$[FC] values. Red genes have a positive $Log_2$[FC] value; blue genes have a negative $Log_2$[FC] value. Light red and light blue genes had a significant adjusted *P* value, represented here on a $Log_{10}$ scale. The top 10 DEGs are shown as enlarged red or blue points.

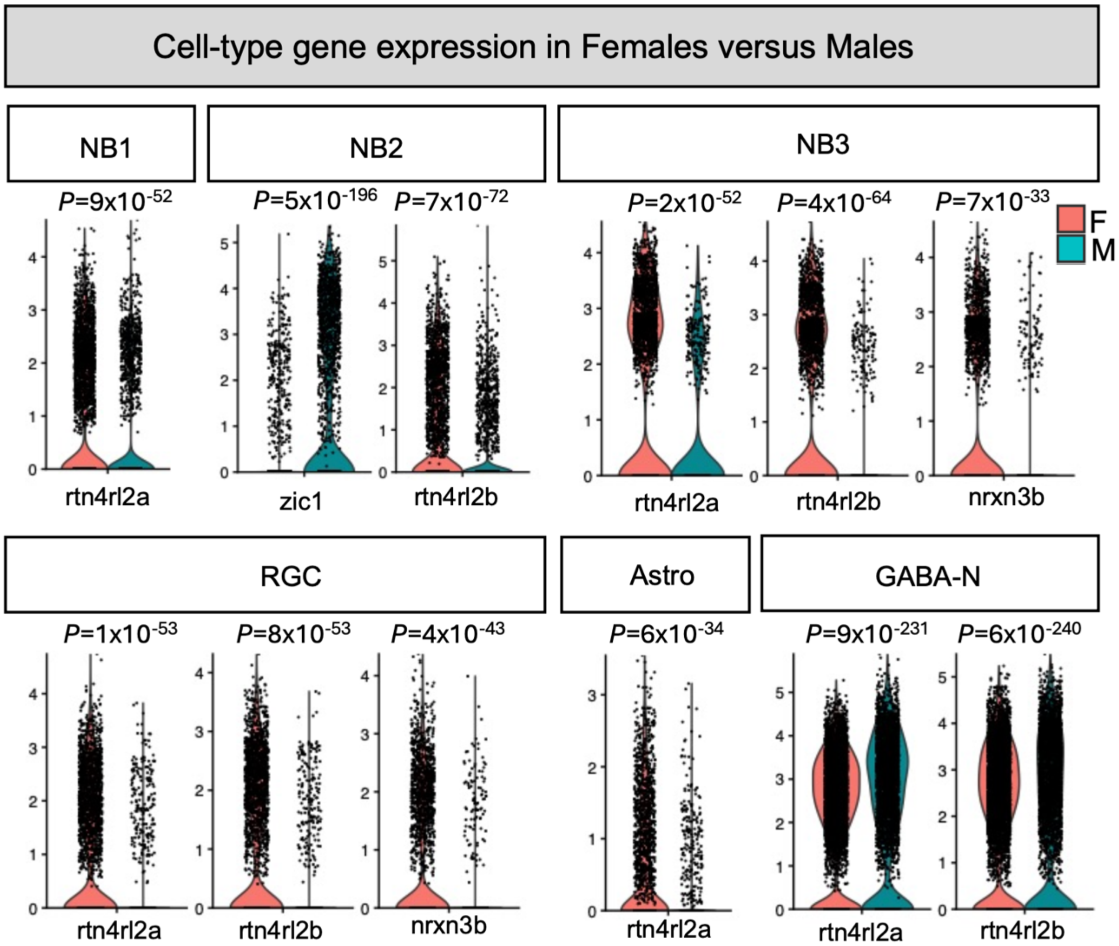


**Figure S8. Sex-dependent differences in cell-type-specific expression of plasticity genes**. Violin plots show gene expression in specific cell types, comparing females and males. The y-axis shows the expression level. Each data point represents a cell. Adjusted p-values are shown directly above each violin plot. The gene for which expression is shown in each violin plot is on the x-axis.

***GO plots** for cell-type gene expression differences for **male versus female** A. burtoni are in a folder titled "Figure S13-Male versus female" in a Box folder titled "Dussenne_etal_2026_Burtoni_hypomap" at this link: https://ucla.box.com/s/eh9azmovdsavicogi9jv75qpp5muplac*

**Figure S9.**

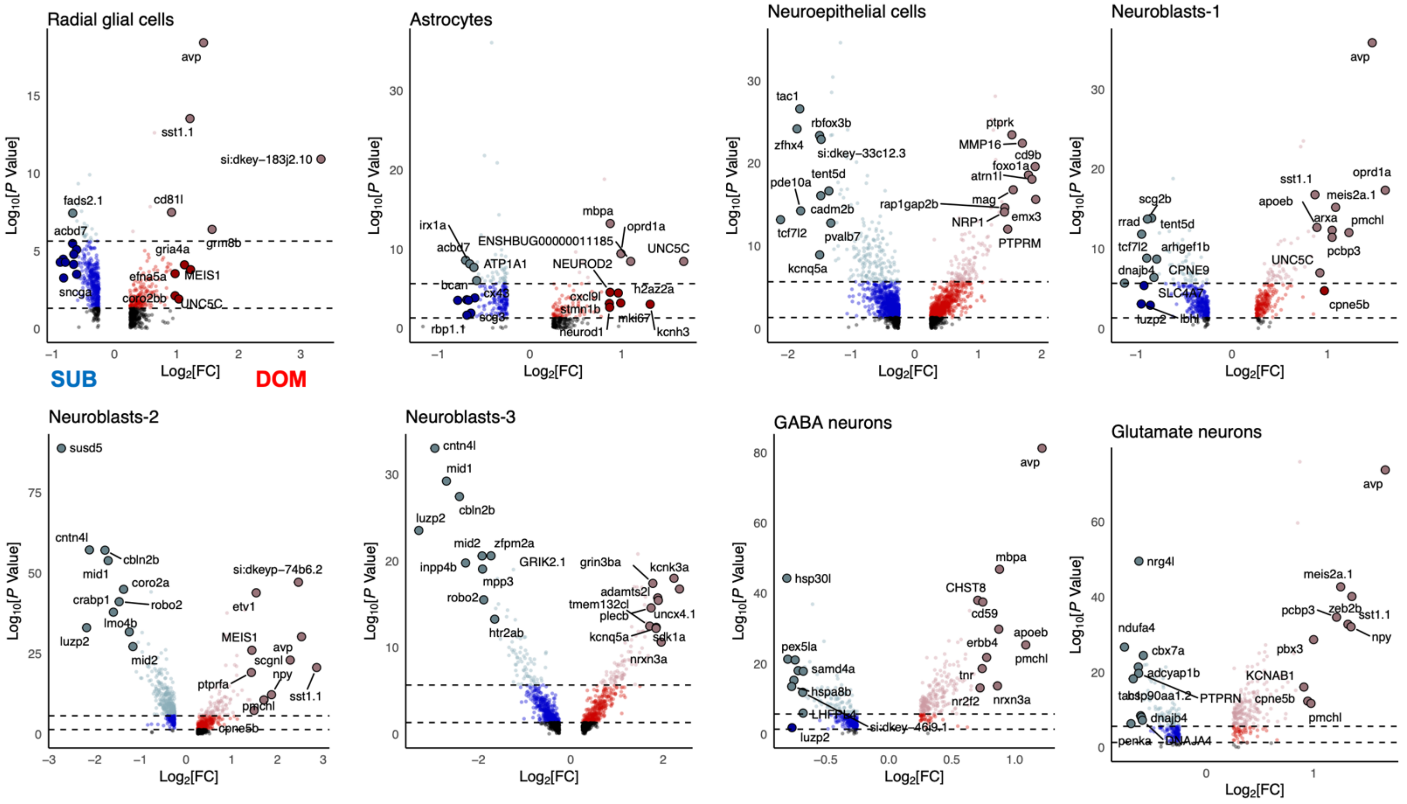


**Figure S10. Status-dependent differences in cell-type-specific expression of plasticity genes**. Violin plots show gene expression in specific cell types, comparing DOM and SUB males. The y-axis shows the expression level. Each data point represents a cell. Adjusted p-values are shown directly above each violin plot. The gene for which expression is shown in each violin plot is on the x-axis.

***GO plots*** *for cell-type gene expression differences for* ***DOM versus SUB*** *A. burtoni are in a folder titled “Figure S13-DOM versus SUB” in a Box folder titled “Dussenne_etal_2026_Burtoni_hypomap” at this link:*
*https://ucla.box.com/s/eh9azmovdsavicogi9jv75qpp5muplac*

**Figure S11**

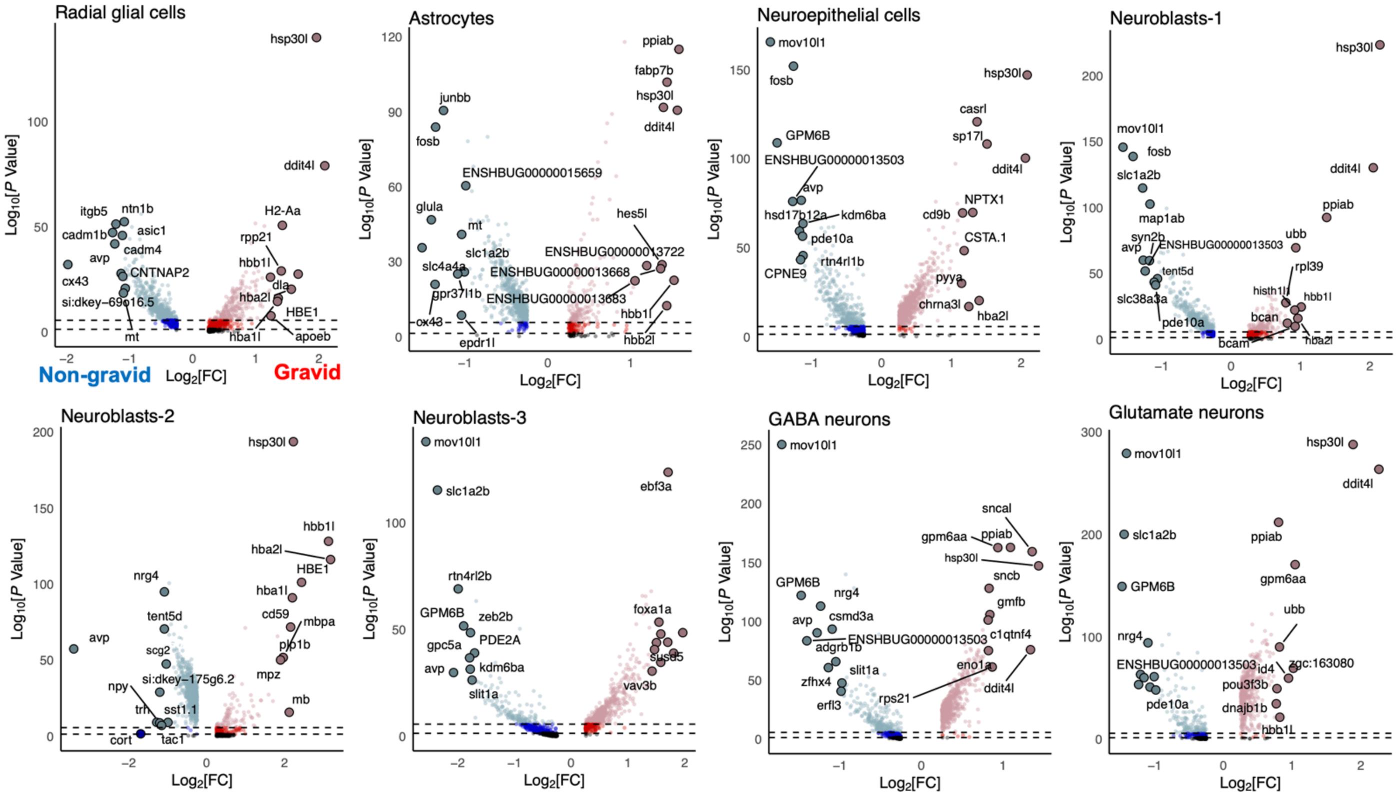


**Figure S12. Female reproductive state-dependent differences in cell-type-specific expression of plasticity genes**. Violin plots show gene expression in specific cell types, comparing gravid and non-gravid females. The y-axis shows the expression level. Each data point represents a cell. Adjusted p-values are shown directly above each violin plot. The gene for which expression is shown in each violin plot is on the x-axis.

***GO plots** for cell-type gene expression differences for **gravid versus non-gravid** female A. burtoni are in a folder titled "Figure S13-Gravid versus non-gravid" in a Box folder titled "Dussenne_etal_2026_Burtoni_hypomap" at this link:*
*https://ucla.box.com/s/eh9azmovdsavicogi9jv75qpp5muplac***]**

**Figure S13.**

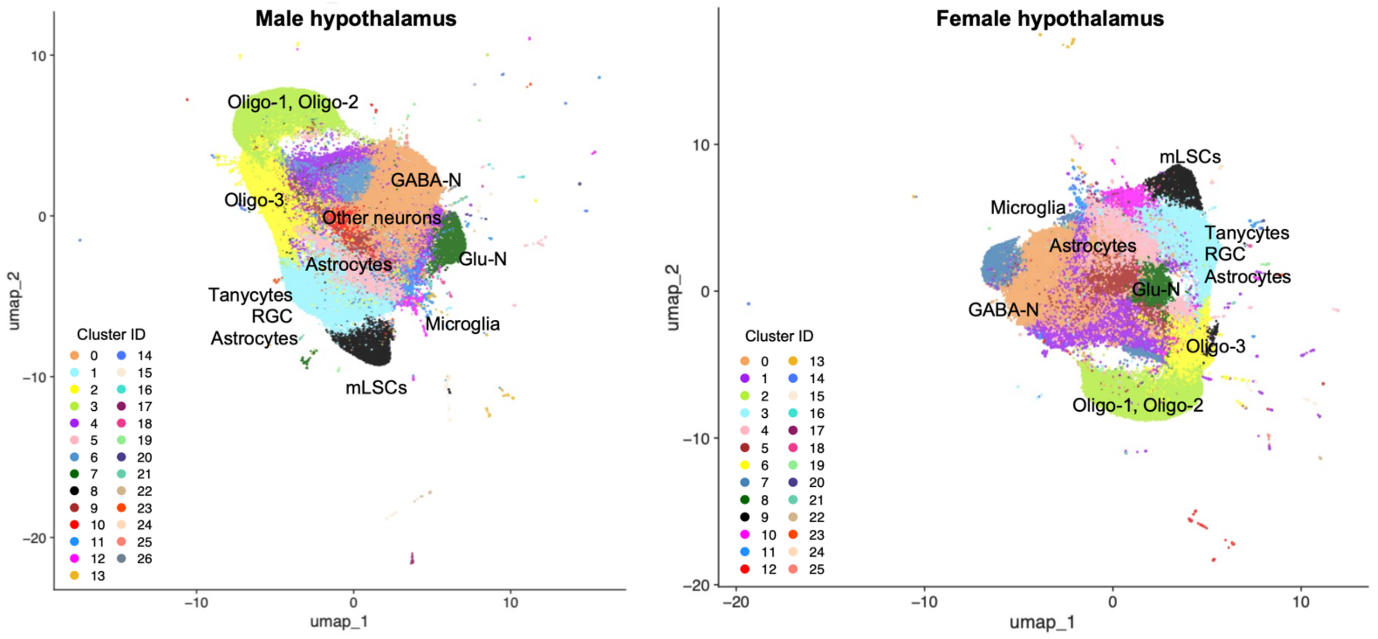


**Figure S14**: **Similarities between the male and female UMAPs.** The main cell types are provided. mLSCs: meningeal lymphatic supporting cells.

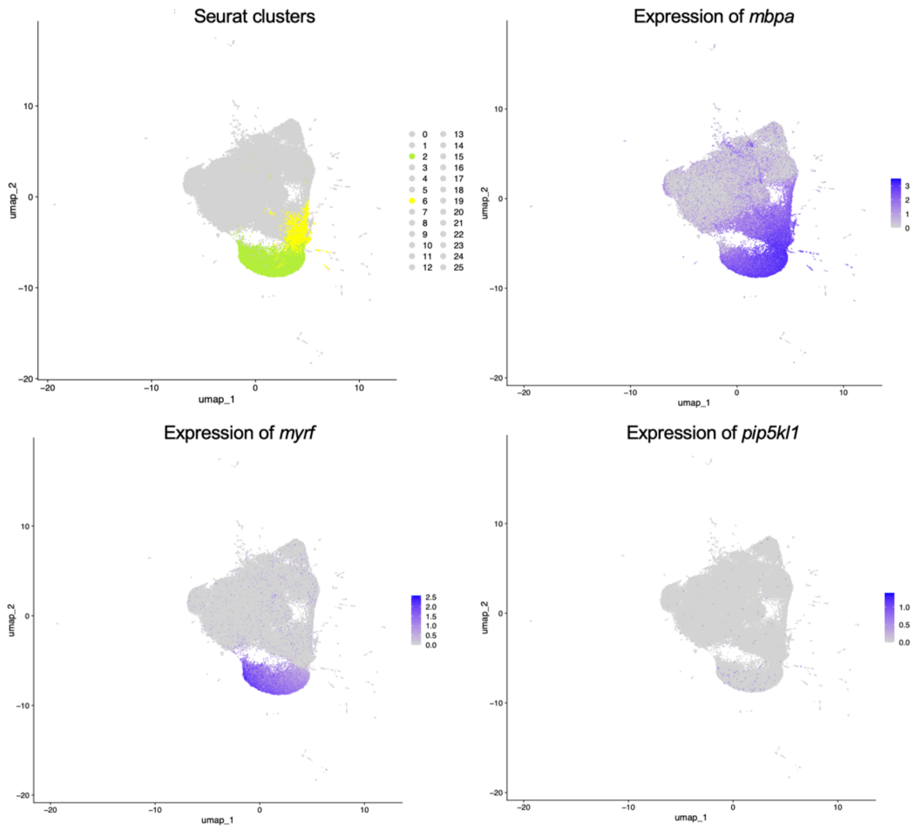


**Figure S15**: **Oligo markers present in the spatial dataset.** UMAPs showing the Oligo-1 and Oligo-2 (F-C2, green), and oligo-3 (F-C6, yellow) cell clusters in the female dataset. Oligo-1 cells present high expression of both *mbpa* and *myrf*. *Pip5kl1* is only expressed in this cluster, possibly labelling Oligo-2 cells. Oligo-3 cells show overexpression of *mbpa*.

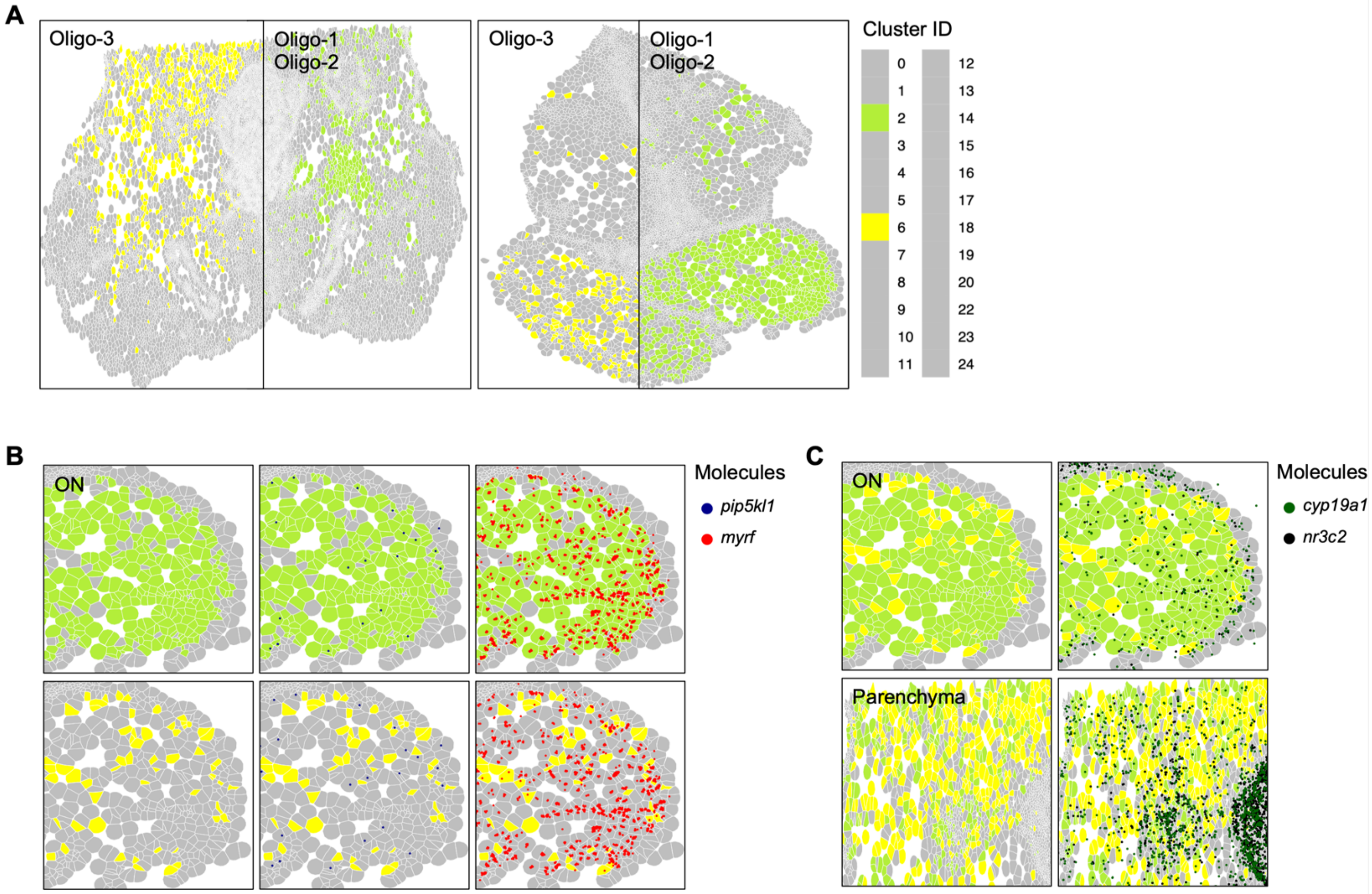


**Figure S16**: **Spatially resolved Oligos and their expression of steroid signaling genes.** Oligodendrocyte clusters in the female brain (F-C2, F-C6), representing Oligo-1, Oligo-2 and Oligo-3 clusters of the scRNA-seq. These cells constitute the majority of the optic nerve (ON) tissue, and are also present in the posterior hypothalamus parenchyma. (A) Oligo-3 cells (F-C6) are less numerous in the parenchyma of the distal hypothalamus compared to Oligo-1 cells (F-C2). On the contrary, Oligo-1 cells are more numerous in the ON than Oligo-3 cells. (B) F-C2 cells strongly express *mbpa*, *myrf*, and *pip5kl1* to a lesser extent. F-C6 is almost only characterized by the overexpression of *mbpa*. (C) Both Oligodendrocyte clusters express *cyp19a1* and *nr3c2* in about 20% of cells.

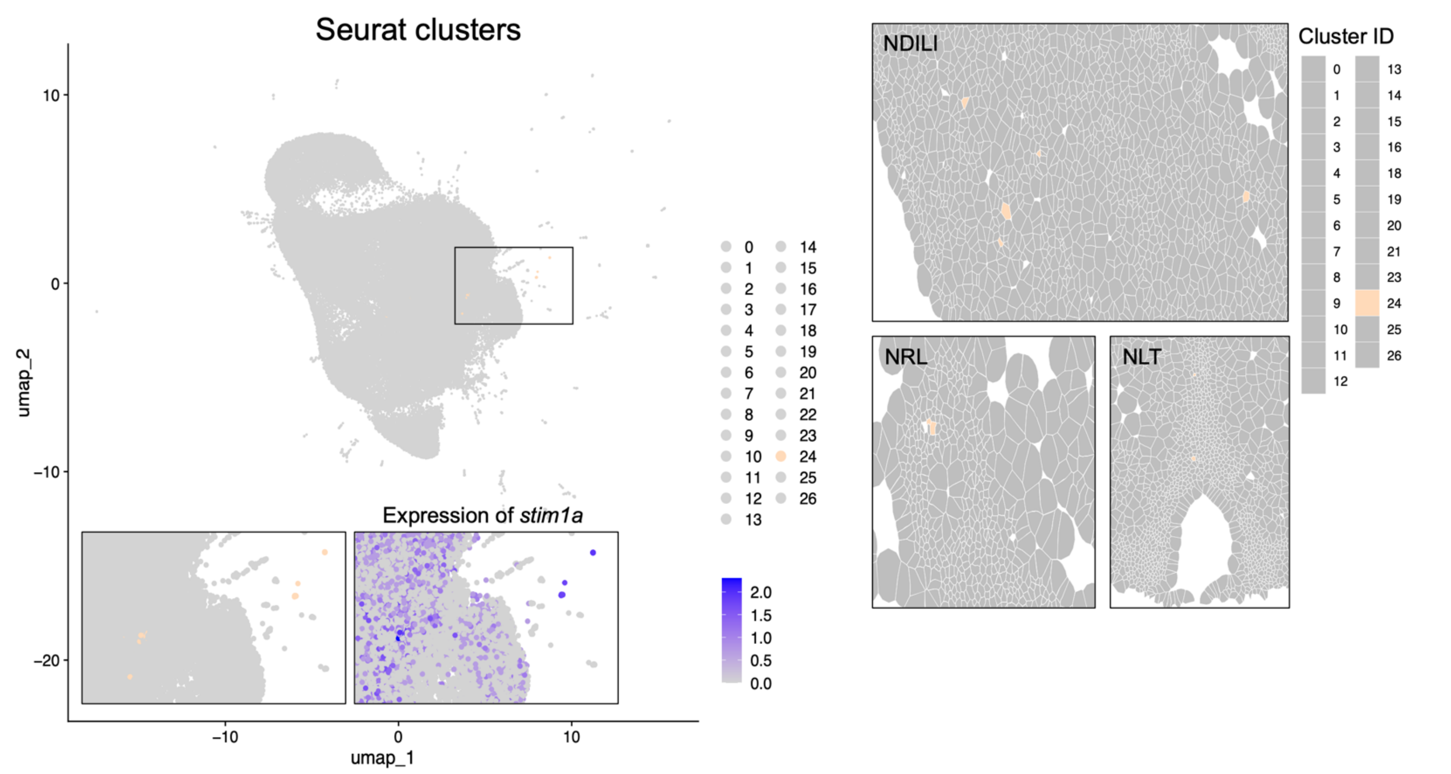


**Figure S17: Identification of a sparsely labeled *stim1a*+ OPC cluster.** UMAP showing the OPC-1 cell cluster (M-C24) in the male hypothalamus, in which *stim1a* is the only marker gene to be significantly expressed in more than 20% of cells. These cells were rare, and located in the NDILI, NRL, NLT and Vv (not shown).

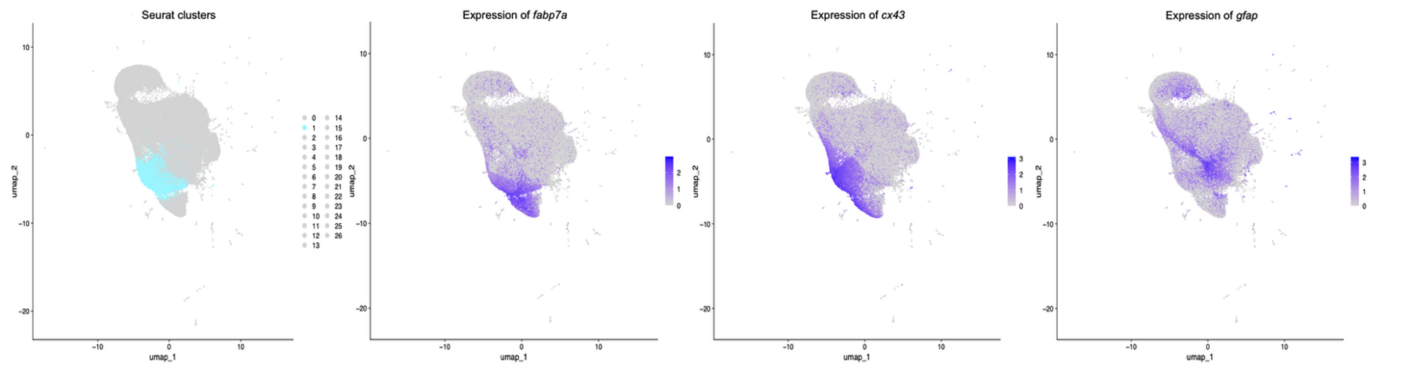


**Figure S18: Glial cell types make up one large cluster.** UMAPs showing M-C1, a cluster proposed to be composed of three cell types: RGC, tanycytes and some astrocytes. These cells overexpress several marker genes, including *fabp7a*, *cx43*, and *gfap* to a smaller extent.

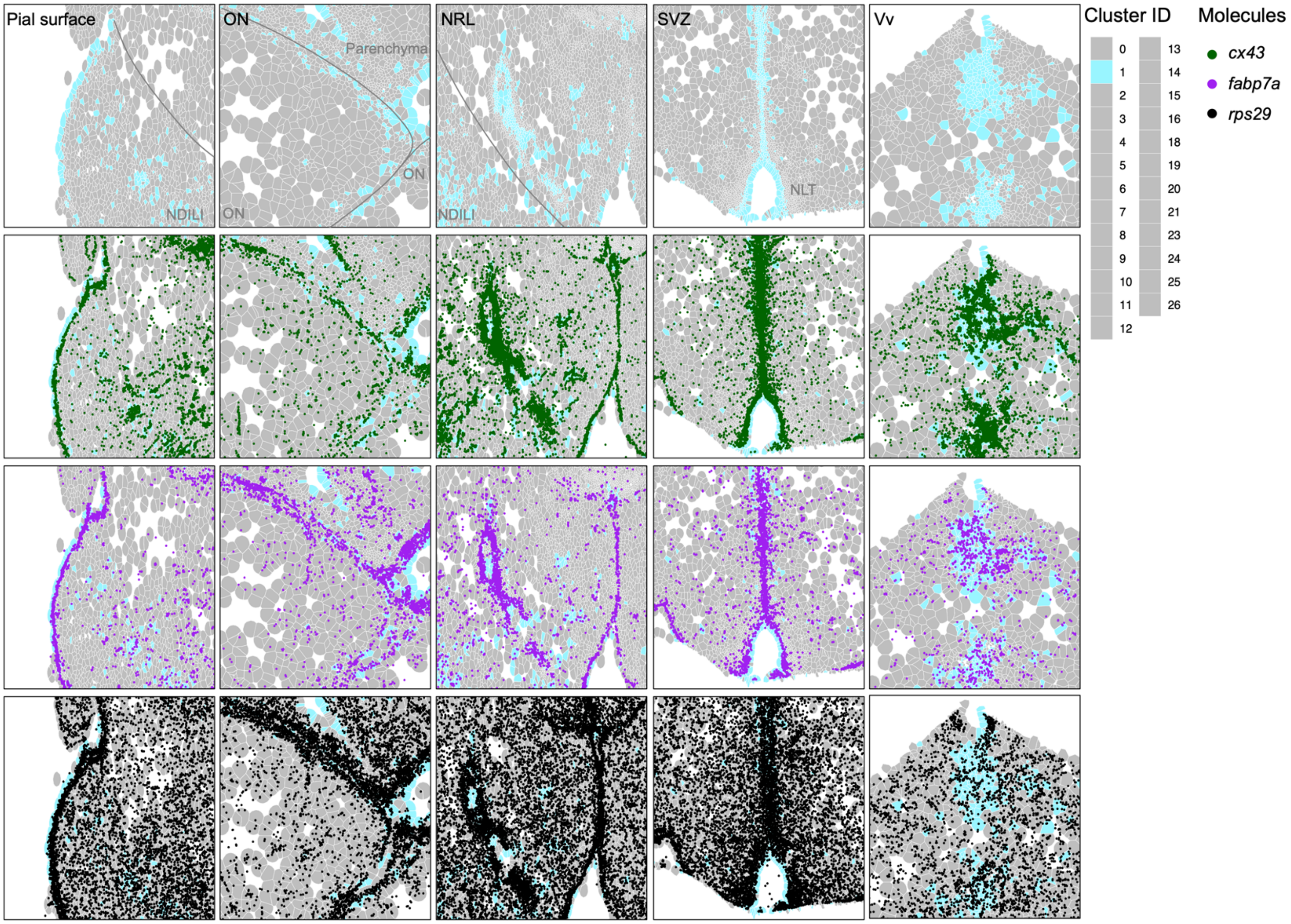


**Figure S19: Spatial distribution of distinct glial cell markers.** Main localization of M-C1 cells in the male hypothalamus (top panel). Lower panels show the localization of transcripts of the three main marker genes in this cluster; *cx43* (dark green), *fabp7a* (purple), *rps29* (black). See Table S2 for abbreviations.

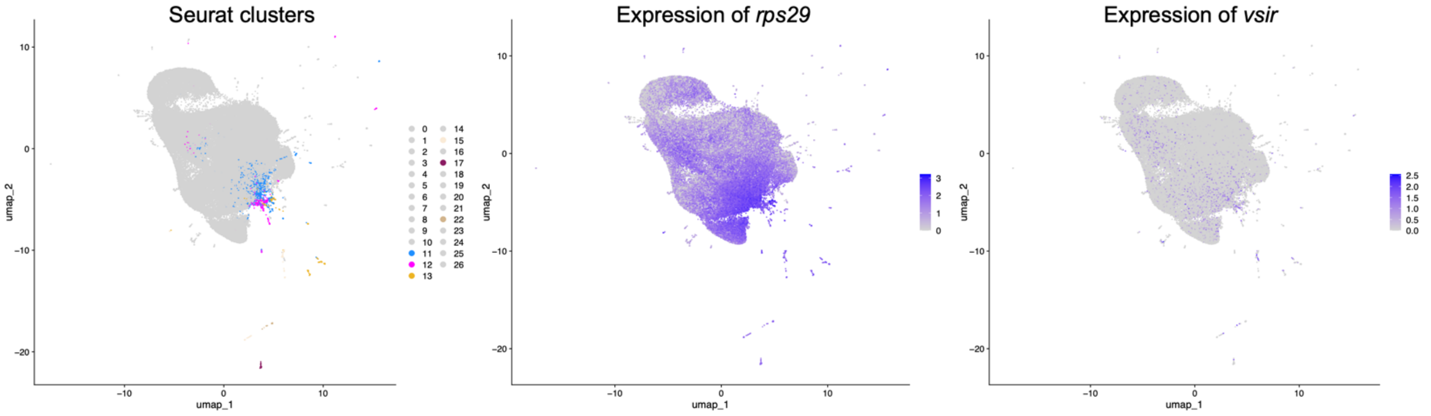


**Figure S20: *Rps29* and *vsir* expression highlights microglial cells in spatial dataset**. UMAP representing microglial clusters in the male dataset (M-C11, 12, 13, 15, 17, 22), characterized by the overexpression of *rps29* and, to a lesser extent, *vsir*.

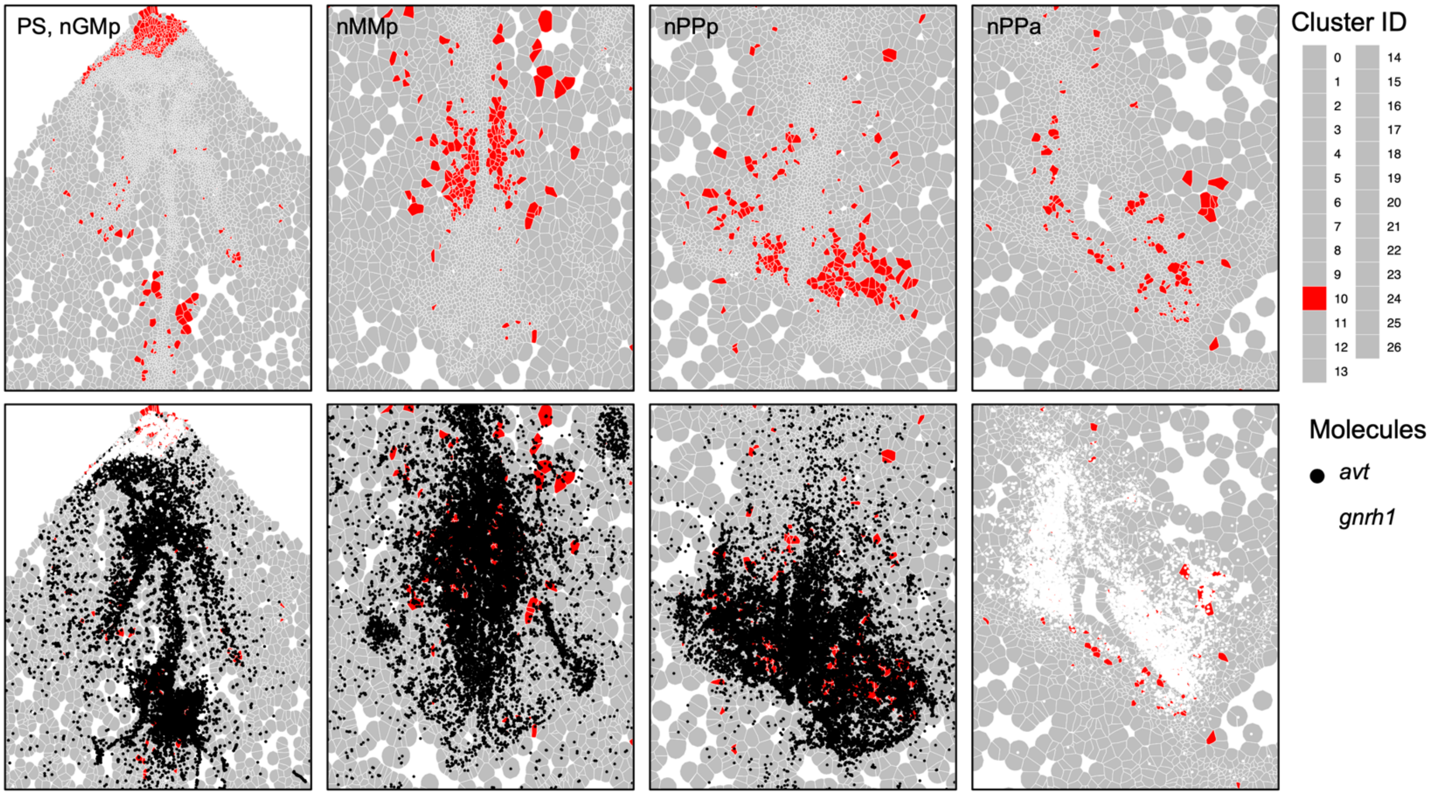


**Figure S21. *Avt* and *gnrh1* overlap with a single cell cluster in the hypothalamus.** M-C10 contained mainly AVT and GnRH1 neurons. These clusters cells are located almost exclusively where *gnrh1* and *avt* transcripts could be found (PS, nGMp, nMMp, nPPa), see Table S2.

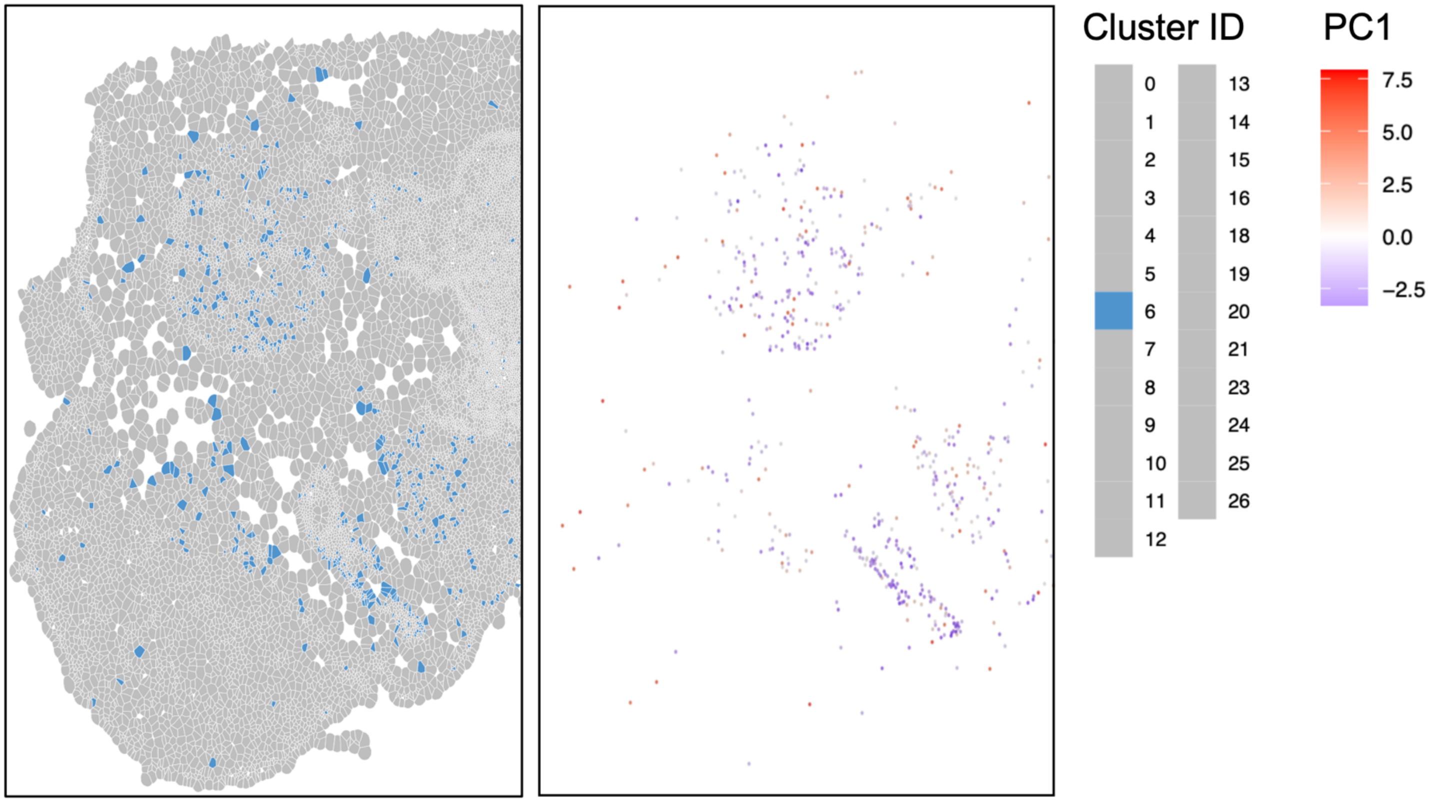


**Figure S22. A spatial gradient was revealed based on microglia and neural gene expression in a male cluster.** Left panel: dispersion of M-C6 cells (blue) across the posterior hypothalamus. Right panel: spatial variation in gene expression within M-C6. The color of each cell depends on its amplitude along PC1, the first principal component. The loadings associated with this PC1 suggest that gradients in gene expression of a few genes drive these gene expression variations. Here, *rps29*, *vsir* and *nrros* are highly expressed in cells colored in red (potential microglial cells), while *grin1a*, *gabbr2*, *cacna2d2a*, and *grin2ab* are highly expressed in blue cells (neurons).

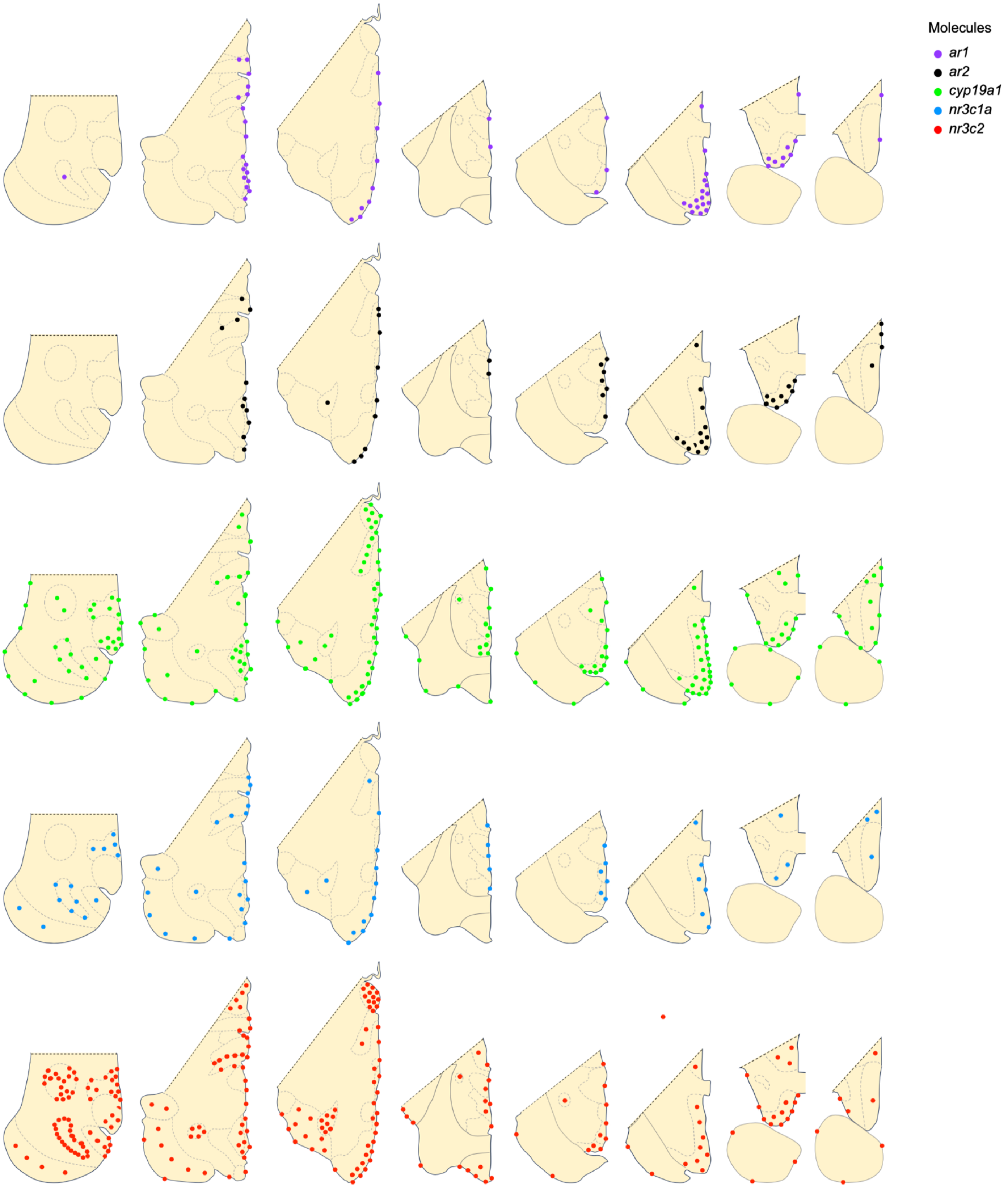
Molecules
ar1
ar2
cyp19a1
nr3c1a
nr3c2

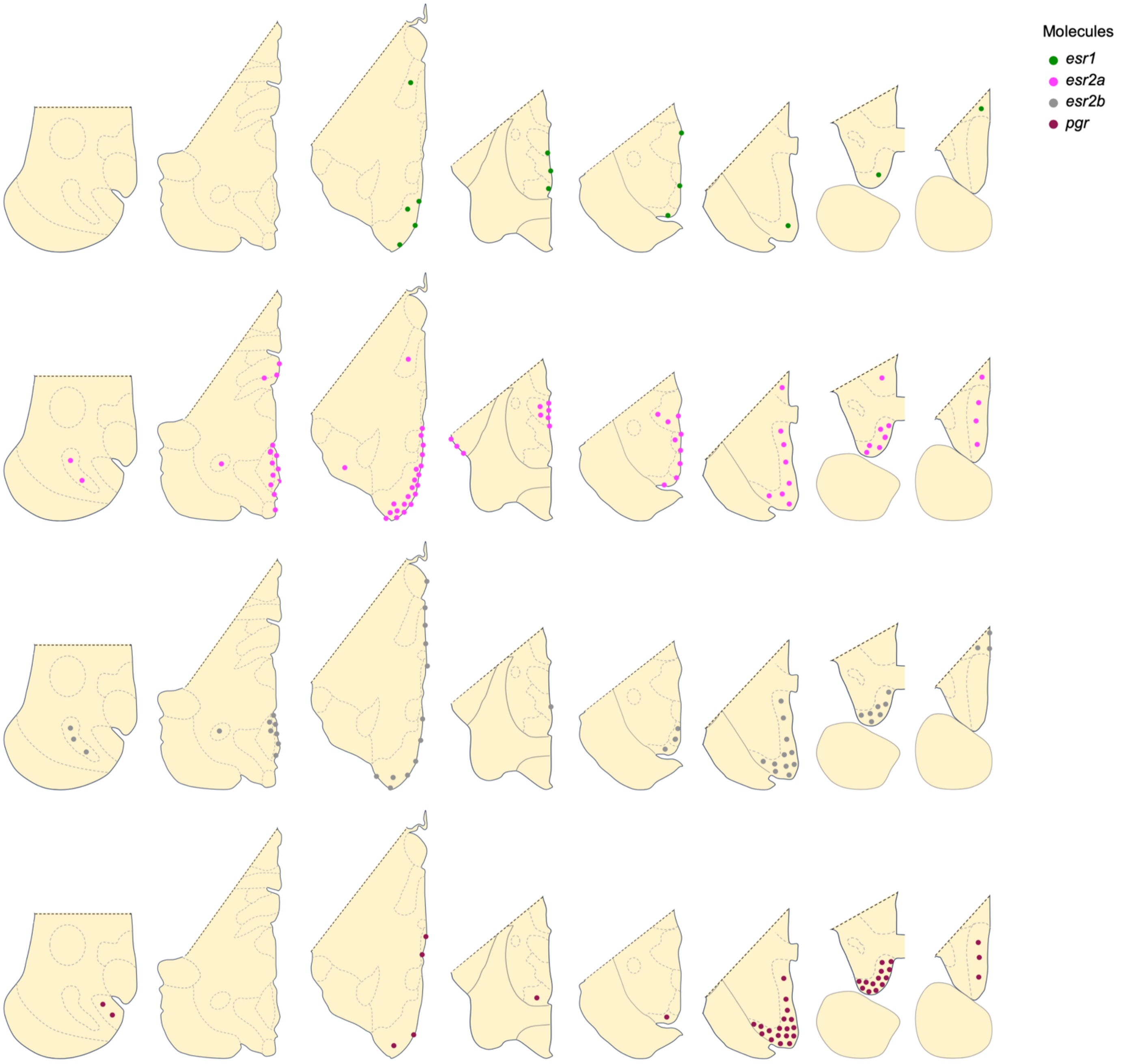


**Figure S23. A spatial catalog of key steroid hormone signaling genes in the *A. burtoni* hypothalamus.** Expression of genes encoding sex steroid receptors, corticosteroid receptors, and aromatase as found in our spatial dataset are schematized, providing a consolidated resource of steroid signaling gene expression in the cichlid brain.

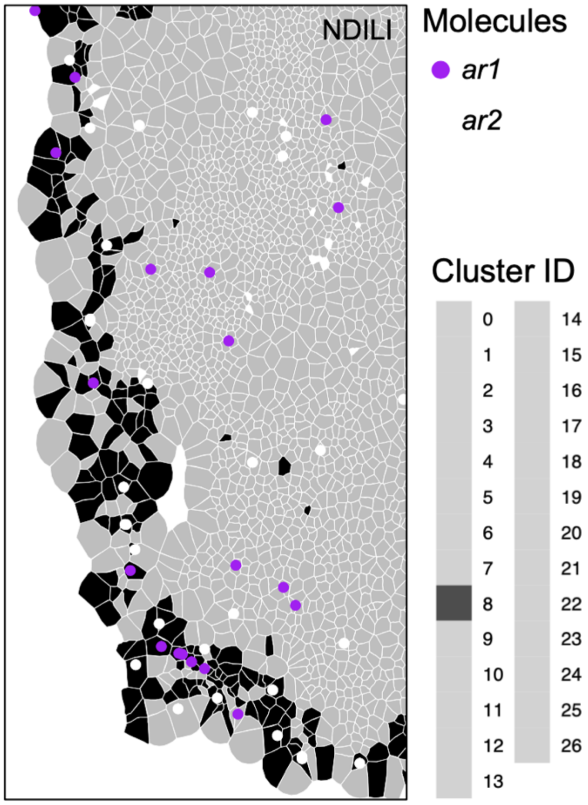


**Figure S24. *Ar* genes are expressed in mLSC cells.** In the male, potential mLSC cells (M-C8, black cells) express *ar1* and *ar2* (purple and white dots, respectively), especially in the posterior hypothalamus. See Table S2 for abbreviations.

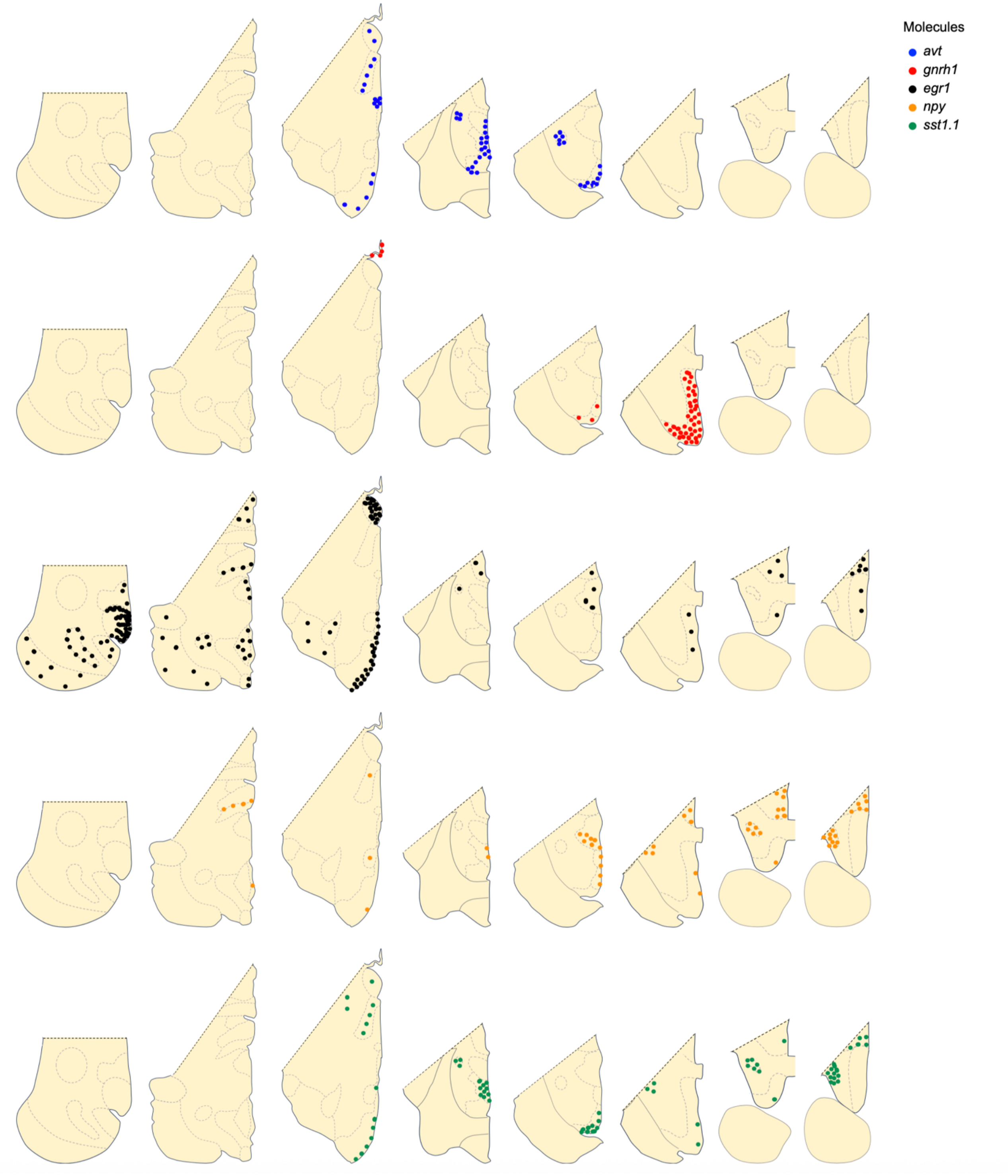


**Figure S25. Distribution of hypothalamic peptidergic neurons and *egr1* in the *A. burtoni* hypothalamus.** Peptide gene expression patterns highlight peptide neuron populations. As in mammals, *npy* and *sst1.1* show some overlap. *Egr1* shows some overlap in expression with *avp* expressing cells.

A

| sgRNA | Direction | Target Sequence |
|---|---|---|
| *sst1.1* crRNA - 1 | reverse | GGAGTCTCTCTCTGCGCGGCGGAGG |
| *sst1.1* crRNA - 2 | reverse | GTGCGCGTGTCGGACCTGTTTGG |

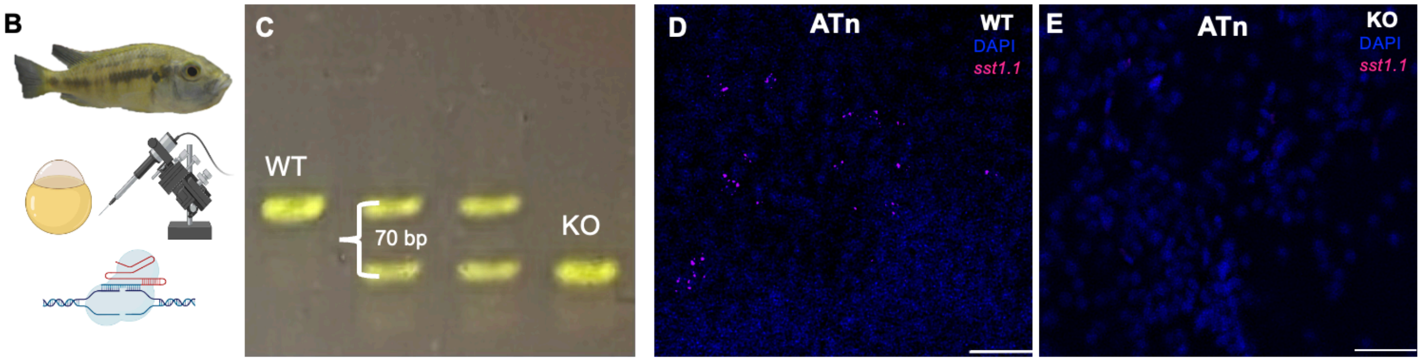


**Figure S26. Generation of *sst1.1* mutant *A. burtoni* using CRISPR/Cas9 gene editing.** A) CRISPR (cr) RNAs were designed against the *sst1.1* gene in CHOPCHOP with the goal of generating large indels. B) Sixty minutes after breeding, fertilized eggs are expelled from the mouthbrooding female and single-cell embryos are injected with the CRISPR/Cas9 gene editing cocktail. C) This produced a 70 bp deletion that was propagated to generate a stable line. Using HCR, *Sst1.1* was detected in (D) WT but not (E) KO brains, providing further support that *sst1.1* KO fish do not make the SST protein. WT=wild-type; KO=knockout; ATn=anterior tuberal nucleus. (D-E) Scale bar=50 μM.

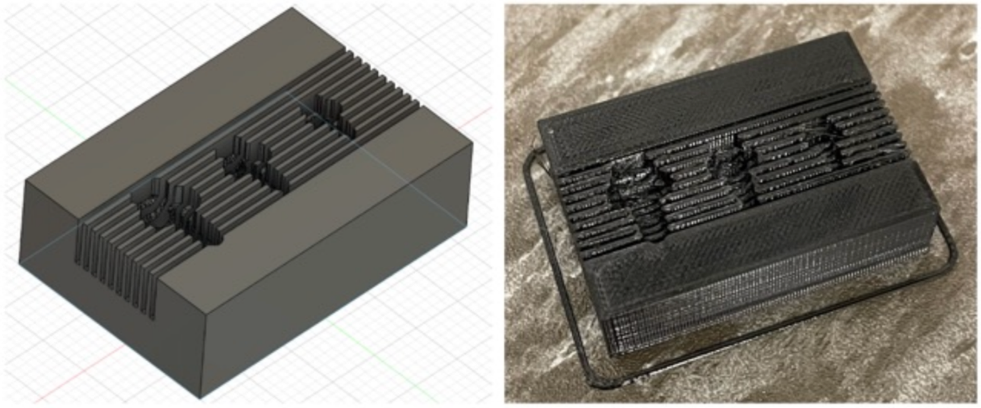

**Figure S27. Custom 3D-printed cichlid brain molds for dissecting brain tissue. [Accessing specs for re-creating/replicating].** Schematic and printed brain molds for dissecting fresh *A. burtoni* brain tissue. Different sized molds allow for best-fit. Fresh razor blades were inserted along the ridges to produced consistently sized brain sections from brains that were fixed in the molds with mow melting-point agarose. Sliced tissue was placed under a dissection microscope to remove parts of the hypothalamus (See Figure 1). Printing was performed with a Prusa 3D printer.
